# Systematic identification of regions where DNA methylation is correlated with transcription refines regulatory logic in normal and tumour tissues

**DOI:** 10.1101/2025.02.03.636201

**Authors:** Richard Heery, Martin H Schaefer

**Affiliations:** Department of Experimental Oncology, IEO European Institute of Oncology IRCCS, Via Adamello 16, 20139, Milan, Italy

## Abstract

DNA methylation at gene promoters is generally considered to be associated with transcriptional repression. However, lack of a clear picture of where promoter methylation is most important for transcriptional regulation has hindered our understanding of this relationship and resulted in the use of a wide variety of arbitrary promoter definitions. We demonstrate here that the use of different promoter definitions can lead to contradictory results between studies of promoter methylation. In response, we have developed Methodical, a computational method that combines RNA-seq and whole genome bisulfite sequencing (WGBS) data to identify genomic regions where DNA methylation is highly correlated with transcriptional activity. We refer to these regions as transcript-proximal methylation-associated regulatory sites (TMRs). We applied Methodical to one normal prostate tissue data set, one prostate tumour dataset and one prostate metastases dataset and characterized the identified TMRs. We show that the region just downstream of the TSS, particularly within the first exon and intron, is the most common location for TMRs and that TMRs are enriched for particular genomic features, chromatin states and transcription factor binding sites. Finally, we demonstrate that the methylation of TMRs is generally strongly correlated with transcription in diverse cancer types and that TMRs are highly subject to altered DNA methylation in cancer.

## Introduction

Gene promoters are broadly defined as the region in the vicinity of transcription start sites (TSS) where RNA polymerase is recruited by transcription factors to initiate transcription. DNA methylation involves the addition of methyl groups to DNA and in mammals generally occurs at the 5’ position of cytosine bases that are followed by a guanine base (CpG sites) (1–3). It is generally accepted that DNA methylation at gene promoters has a repressive effect on transcription (3–5). Two main models have been put forward to describe how DNA methylation could mediate this repression: The first proposes that transcription factor binding is blocked indirectly via recruitment of methylated DNA-specific proteins which in turn recruit co-repressors that silence transcription, while the second proposes that DNA methylation directly blocks transcription factor binding (4).

However, genome-wide studies have reported fairly weak correlations in general between promoter DNA methylation and gene expression (6–8). Additionally, exceptions to the general paradigm of gene silencing by DNA methylation have been reported where DNA methylation has been found to be positively associated with the expression of certain genes (9, 10). Thus, the relationship between DNA methylation and transcriptional activity seems to be more nuanced and context-dependent than suggested by the conventional model. Moreover, DNA methylation at regions downstream of the TSS, particularly the first intron and exon, have been recognized DNA methylation to be associated with transcriptional silencing (11, 12). Additionally, variably methylated regions outside promoter regions were found to often display stronger correlations with gene expression than promoter regions when studying single cells (13).

When studying promoter DNA methylation, typically CpG methylation values have been averaged over the extent of the designated promoter region and linear models then fit to gene expression values. Several recent studies have shown that this simple approach often fails to accurately capture the relationship between DNA methylation and transcription, with more sophisticated models incorporating distal regions and considering methylation at a higher resolution providing far more accurate predictions of transcriptional activity. In particular, a probabilistic machine learning approach which extracted higher order methylation features produced a very powerful predictor of gene expression (14). Alternative machine learning approaches have also provided further insight into the association between DNA methylation and transcription, highlighting the importance of the region just downstream of the TSS as well as distal regions (15–17). Instead of directly predicting gene expression, another approach sought to predict promoter activity indirectly by using the enrichment of H3K4me3 and H3K27ac around the TSS as a proxy and predicting their levels using DNA methylation. The predicted levels of these marks in turn were found to be relatively well correlated with transcriptional activity as measured by RNA-seq (18).

Another major limitation of previous studies of promoter DNA methylation has been the lack of agreement on the extent of promoters around TSS, resulting in the use of a wide variety of completely arbitrary promoter definitions. These different definitions have varied from hundreds to thousands of base pairs in length and also differed in the proportion of the promoter sequence located downstream of the TSS relative to upstream (19–23). Indeed, it is difficult to find two studies using precisely the same promoter definition.

Variation in choice of promoter definition could obviously lead to inconsistent results between studies of promoter DNA methylation, including those aiming to identify promoters affected by DNA methylation change in cancer. Additionally, alternative promoter definitions could lead to the calculation of different correlation values between promoter methylation and transcript expression, obscuring our understanding of this relationship and also the recognition of which methylation changes in cancer are associated with corresponding transcriptional changes.

Most large-scale studies of DNA methylation over the last decade and a half have utilized the Illumina methylation microarrays due to their relative cost-effectiveness. This includes projects studying molecular alterations in diverse cancer types such as TCGA (24) and TARGET (25). However, these microarrays measure DNA methylation only at a small percentage of the over 29.4 million CpG sites present in the human reference genome. The Infinium HumanMethylation450 array measures methylation at about 450, 000 CpG sites, 1.5% of the total number present in the genome and mostly targets CpGs located in CpG islands and promoters. The most recent generation of Illumina methylation microarray, the Infinium MethylationEPIC v2.0 array, targets over 935, 000 CpG sites, about 3% of the total number, and has expanded the coverage of other regions compared to the HumanMethylation450 array, particularly of enhancers. It still however only covers about 3% of human CpG sites and the targeted CpG sites are still highly enriched for certain genomic regions, such as CpG islands (26). Thus, most attempts to study the relationship between DNA methylation and transcription across the genome have been limited to a small proportion of CpG sites and biased towards certain genomic contexts (27, 28).

The publication over the last few years of datasets with whole genome bisulfite sequencing (WGBS) and RNA-seq data for large numbers of human samples provide the opportunity to search for regions where DNA methylation is highly correlated with transcription in an unbiased manner. Thus, we developed Methodical, an algorithm which systematically identifies regions displaying such correlations. Two of the largest datasets to date have been for prostate cancer: one for prostate tumours and matching normal prostate samples (21) and another for prostate metastases (29). We divided the prostate tumour and matching normal prostate dataset into two separate datasets for prostate tumours and normal prostate samples in order to study the relationship between DNA methylation and transcription in cancer and normal tissues separately. We applied Methodical to the three different datasets and subsequently characterized the identified regions, revealing novel insights into the relationship between DNA methylation and transcription in both normal prostate tissue and prostate cancer.

## Materials and methods

### Datasets

We used three different datasets to identify TMRs: one for normal prostate with 126 samples, one for prostate tumours with 126 samples and one for prostate metastases with 99 samples. The normal prostate and prostate tumour samples come from the CPGEA project (21) and the prostate metastasis samples come from the MCRPC project (29).

### Location of TSS and transcript-encoding regions

The location of TSS and the associated transcript-encoding regions for all protein-coding transcripts were obtained from the Gencode (https://ftp.ebi.ac.uk/pub/databases/gencode/Gencode_human/release_38/ <u>gencode.v38.annotation.gtf.gz</u>). The genomic location of the first base of each transcript was designated as a TSS. To identify a subset of high-confidence TSS from among these, we used data produced by cap analysis of gene expression (CAGE), a technique to profile the 5’ ends of mRNA molecules(30), from the FANTOM5 project (31, 32).

We downloaded CAGE data for human prostate (http://fantom.gsc.riken.jp/5/datafiles/latest/basic/human.tissue.hCAGE/prostate%252c%2520adult%252c%2520pool1.CNhs10628.10022-101D4.hg19.ctss.bed.gz) and filtered for TSS supported by at least 10 CAGE tags. This gave us a set of 17, 071 high-confidence TSS for 10, 027 different genes. We used this set of TSS for all analyses unless stated otherwise.

### Transcript Expression

Transcript sequences were downloaded from Gencode (https://ftp.ebi.ac.uk/pub/databases/gencode/Gencode_human/release_38/ gencode.v38.transcripts.fa.gz). The expression of transcripts was quantified from paired end RNA-Seq FASTQ files using Kallisto (version 0.46.1) (33) with 100 bootstrap samples. Transcript counts were then normalized using the median of ratios approach from DESeq2 (version 1.38.1) (34).

### DNA Methylation-Transcription Correlations and Methylation of Genomic Regions

All correlation values mentioned are Spearman correlation values. Statistical significance of correlation values was calculated by using the pt() function in R with the t-statistics derived from the correlation values. When calculating the methylation values of genomic regions, the mean of the methylation values of all CpG sites overlapping them was used. Differential methylation for genomic regions was tested using the diff_methylsig() function from the methylSig R package (35) (version 1.10.0) using the number of methylated reads and total number of WGBS reads overlapping each region.

### Gene Set Overrepresentation Analysis

MSigDB Hallmark pathways and KEGG pathways were obtained from MSigDB (36, 37) version 7.5 using the msigdb R package (version 1.6.0). Pathway overrepresentation among TMR-associated genes was performed by taking all genes associated with a given group of TMRs and testing the significance of their overlap with gene sets using the one-sided version of Fisher’s exact test using all protein-coding genes as the background. p-values were adjusted using the Benjamini-Hochberg procedure.

### Genomic Features, Chromatin States and Repeats

The location of introns and exons for protein-coding genes were obtained from the Gencode 38 comprehensive gene annotation. The locations of CpG islands, shores shelves and open sea regions were <u>obtained</u> using the annotatr R package (38) (version 1.24.0). Predicted promoter regions, predicted enhancer regions, open chromatin regions and CTCF binding sites were downloaded from Ensembl version 109 using the biomaRt R package (39) (version 2.54.0).

Chromatin states for prostate tissue were downloaded from https://download.cncb.ac.cn/OMIX/OMIX237/OMIX237-64-02.zip and lifted over to hg38 using the rtracklayer R package (40) (version 1.58.0). The location of repeat sequences were downloaded from the UCSC genome browser (https://hgdownload.soe.ucsc.edu/goldenPath/hg38/bigZips/hg38.fa.out.gz).

### Transcriptional Regulator Binding Site Enrichment

The locations of binding sites for transcriptional regulators were obtained from ReMap2022 (41). The BED file for non-redundant peaks in human for hg38 was downloaded from https://remap.univ-amu.fr/storage/remap2022/hg38/MACS2/remap2022_nr_macs2_hg38_v1_0.bed.gz and filtered for any binding sites identified in primary prostate tissue or prostate tumours or else cell lines derived from prostate tissue or prostate cancer (22Rv1, DU145, DuCAP, LAPC-4, LNCaP, LNCaP-95, LNCaP-abl, LNCaP-C4-2, LNCaP-C4-2B, LNCaP-FGC, MDA-PCa-2b, PC-3, RWPE-1, RWPE-2, VCaP, VCaP-LTAD and WPMY-1). There were 77 different transcriptional regulators which had binding sites in prostate tissue, tumours or cell lines.

The enrichment of binding sites among TMR groups was tested by comparing the proportion of CpG sites within TMRs which overlap binding sites with that of all CpGs within the search region for TMRs (transcript-encoding regions with 5 kb added upstream and downstream) using a two-sided chi-squared test. p-values were adjusted using the Benjamini-Hochberg procedure.

### Methodical Algorithm

Spearman correlation values are calculated between the expression of a given transcript and the methylation values of all CpG sites within a specified region surrounding the TSS. We used the entire transcript-encoding region plus 5 kb upstream of the TSS and 5 kb downstream of the transcription end site (TES) of each transcript. The significance of these correlations is inferred and the resulting p-values are corrected for multiple testing using the Benjamini-Hochberg procedure. These corrected p-values are then transformed by taking their logarithm to the base 10 and multiplying by -1 if the correlations are positive, giving what we term the Methodical scores for the correlations. Thus, positive correlations have positive Methodical scores and negative correlations have negative Methodical scores, with the magnitude of the scores determined by the statistical significance of the correlations.

Smoothing of data across CpG sites has previously been employed to reduce noise in WGBS data (42) and Methodical adapts that approach by smoothing scores using an exponential moving average. The moving average employs a window which is centred on a given CpG site and includes an equal number of flanking CpGs upstream and downstream of this central CpG. For example, with five flanking CpGs, there would be a window size of 11 consisting of the central CpG, five CpGs upstream and five CpGs downstream. Weights decay geometrically and symmetrically moving away from the central CpG across the other CpGs in the window using a specified smoothing factor.

Two symmetrical significance thresholds are used to identify negative and positive TMRs: A Methodical score of log_10_(0.05) is used to identify negative TMRs and a Methodical score of -log_10_(0.05) to identify positive TMRs. Wherever the smoothed Methodical scores exceeds one of these thresholds, we group all CpGs within that region as a TMR of the corresponding direction.

We assessed different combinations of the smoothing factor for the exponential moving average and the number of flanking CpGs used to construct windows. We then evaluated the TMRs identified in the metastasis samples by calculating the correlations between methylation of the identified TMRs and expression of their associated transcript in prostate tumour samples. This approach allowed us to gauge the reproducibility of TMRs across different datasets. We found that a smoothing factor of 0.75 and windows constructed using 10 flanking CpGs resulted in a good trade-off between the number of TMRs identified and their reproducibility (Supplementary Figure 1A and B).

### Refinement of Methodical

We investigated if the number of CpGs contained by TMRs was associated with the correlation between their methylation level and transcription of their associated transcript, which we will henceforth refer to just as the TMR correlation values. We discovered that there was a significant correlation between the number of CpGs TMRs contained and the strength of their correlation values (Supplementary Figure 2). We thus decided to filter TMRs for those containing at least 5 CpG sites.

We discovered a huge number of positive TMRs in the prostate tumour and metastasis samples, particularly beyond a few kb from the TSS where they became far more common than negative TMRs (Supplementary Figure 3). Attempts to correct for tumour purity did not change this. We thus hypothesized that the huge number of positive TMRs in cancer samples were possibly related to difficulty mapping WGBS reads to repetitive regions, leading to alignment ambiguity and inaccurate methylation calls in these regions. Indeed, when we evaluated the overlap of repetitive elements, we found that the proportion of TMRs identified in tumour and metastasis samples overlapping repeats increased steadily as the search region was enlarged from 5KB to 50KB and 500KB (Supplementary Figure 4). This phenomenon was particularly pronounced for positive TMRs and the LINE class of repeats, with over 30% of positive TMRs identified in tumour samples overlapping LINE elements when the search region was +/- 500 KB from the TSS.

We thus decided to discard all TMRs which even partially overlapped repeats as we felt we could not be confident in their validity. We reexamined the distribution of TMRs around the TSS after discarding repeat-overlapping TMRs and found that the vast majority of TMRs located at a large distance from the TSS when using large search windows were removed, leaving a clear large peak of negative TMRs and a smaller peak of positive TMRs close to the TSS (Supplementary Figure 5).

We finally evaluated how the number of TMRs identified varies with the number of samples by counting the number of TMRs identified when running Methodical with differently sized subsets of normal prostate samples. We found that with less than 60 samples, few TMRs are identified (Supplementary Figure 6) hinting that a minimum cohort size is required for a useful application of Methodical.

## Results

### Variable Promoter Definitions Lead to Inconsistent Differential Methylation Results

Studies of promoter DNA methylation have employed a variety of different promoter definitions. Figure 1A shows five different promoter definitions used in five different published studies of DNA methylation in cancer (A(19), B(20), C(21), D(22), E(23)). Use of such different promoter definitions could obviously result in different promoter methylation levels being calculated for the same gene or transcript, potentially having a drastic impact on differential methylation results. To investigate this potential problem thoroughly, we decided to compare the differential promoter methylation results obtained using different promoter definitions with the same data set. Thus, we evaluated differential promoter methylation testing in prostate tumour samples compared to matching normal prostate samples using each of the five promoter definitions from Figure 1A with a prostate cancer WGBS dataset (21).

**Figure 1:**
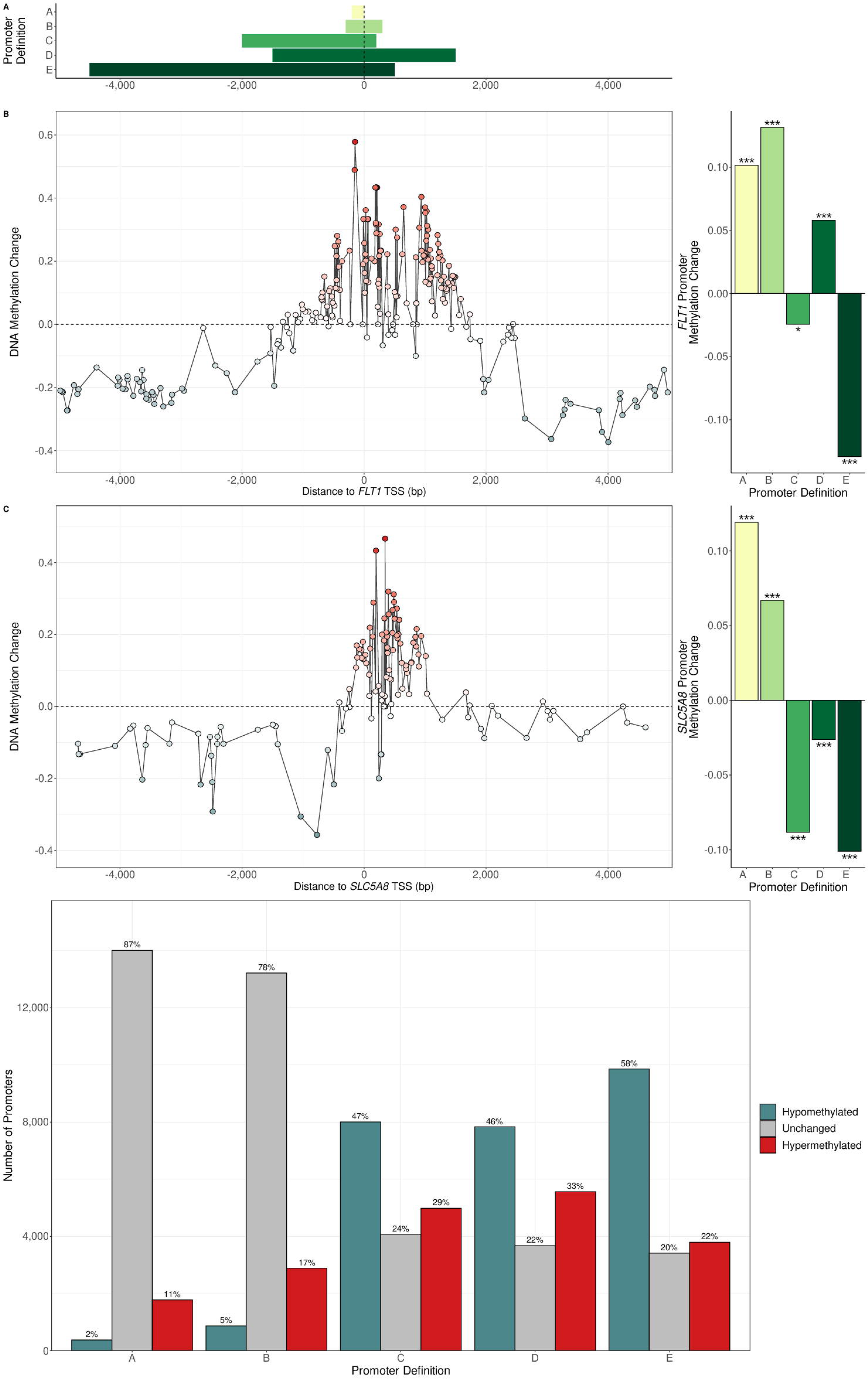
(A) The location of five different promoter definitions relative to the TSS from five different studies of DNA methylation. (B and C) The effect on the choice of promoter definition on differential promoter methylation analysis for *FLT1* (using the TSS associated with the canonical transcript ENST00000282397) and *SLC5A8* (using the TSS associated with the canonical transcript ENST00000536262). In the left-hand plots, the x-axes show distance of CpG sites upstream and downstream of the TSS in base pairs and the y-axes show mean change of methylation in prostate tumour samples compared to normal prostate samples. The right-hand plots show the promoter methylation change calculated using different promoter definitions by averaging the methylation values across CpG sites overlapping each definition. Stars indicate the level of statistical significance for methylation change using a beta-binomial test (*** indicates a p-value < 0.001, ** indicates a p-value < 0.01 and * indicates a p-value < 0.05). With definitions A, B and D the *FLT1* promoter is hypermethylated, while using definitions C and E it is hypomethylated. With definitions A and B, the *SLC5A8* promoter is hypermethylated while for definitions C, D and E it is hypermethylated. (D) The number of significantly hypermethylated and hypomethylated promoters when using the different promoter definitions in the same prostate cancer dataset. Figures above the bars give percentages out of the total of all promoters tested. Promoter definition choice substantially affects the number of differentially methylated promoters discovered, with longer definitions resulting in greater numbers of differentially methylated promoters, especially the number of hypomethylated promoters.

Strikingly, we observed that different promoter definitions can lead to completely opposing results for the same TSS. Figure 1B and 1C show the effect of the choice of promoter definition on differential promoter methylation for the genes *FLT1*, which encodes the vascular endothelial growth factor receptor 1 and which has been reported as hypermethylated in prostate cancer (43), and *SLC5A8,* a candidate tumour suppressor gene also reported as hypermethylated in prostate cancer (44). We found that their promoters were identified as hypermethylated when using one set of promoter definitions but hypomethylated when using another set. We noted that there can be more than a thousand TSS where a given pair of different promoter definitions lead to such contradictory results (Supplementary Figure 7).

We also observed that the choice of promoter definition drastically impacted the number of differentially methylated promoters found, with the wider definitions C, D and E resulting in several times the number of differentially methylated promoters than the narrower definitions A and B (Figure 1D). The number of hypomethylated promoters identified was the most affected, likely reflecting the tendency of genomic regions further away from TSS to lose methylation in cancer. We found that among all promoters identified as differentially methylated with at least one of the five promoter definitions, only a tiny minority were common to all five definitions, with a large proportion exclusively identified with a single one of the definitions (Supplementary Figure 8A and B). For example, 1, 268 promoters were found to be hypermethylated only when using promoter definition D, but not with any of the other definitions, while 2, 071 promoters were found to be hypomethylated only with promoter definition E, but not with any of the others.

### Fixed Promoter Definitions Display Poor Correlations between Methylation and Transcriptional Activity

Promoter DNA methylation is generally of interest as it is assumed that it is associated with transcription of the associated gene. Given the substantial influence of choice of promoter definition on the calculation of promoter methylation levels, it seems obvious that promoter definition could also have a large influence on the calculation of correlation values between promoter methylation and transcript expression. To investigate this, we decided to examine how the correlation values between DNA methylation and transcriptional activity varied with distance of CpG sites from the TSS and if one promoter definition tended to capture the regions with the strongest correlations.

We discovered that there could be substantial variation in these correlation values, with groups of CpG sites with strongly negative or positive correlations found interspersed among CpGs with weak correlations and that fixed promoter definitions often failed to capture this complexity. Figure 2B and 2C show the correlations values around *FOXD1* and *PACSIN3* in prostate metastases as examples. Despite the presence of CpG sites near the TSS where DNA methylation levels are relatively strongly correlated with transcription, fixed promoter definitions often resulted in poor correlations because they either failed to include these CpG sites, included many CpG sites where DNA methylation is not correlated with transcription or overlapped CpG sites with both negative and positive correlations with transcription.

**Figure 2:**
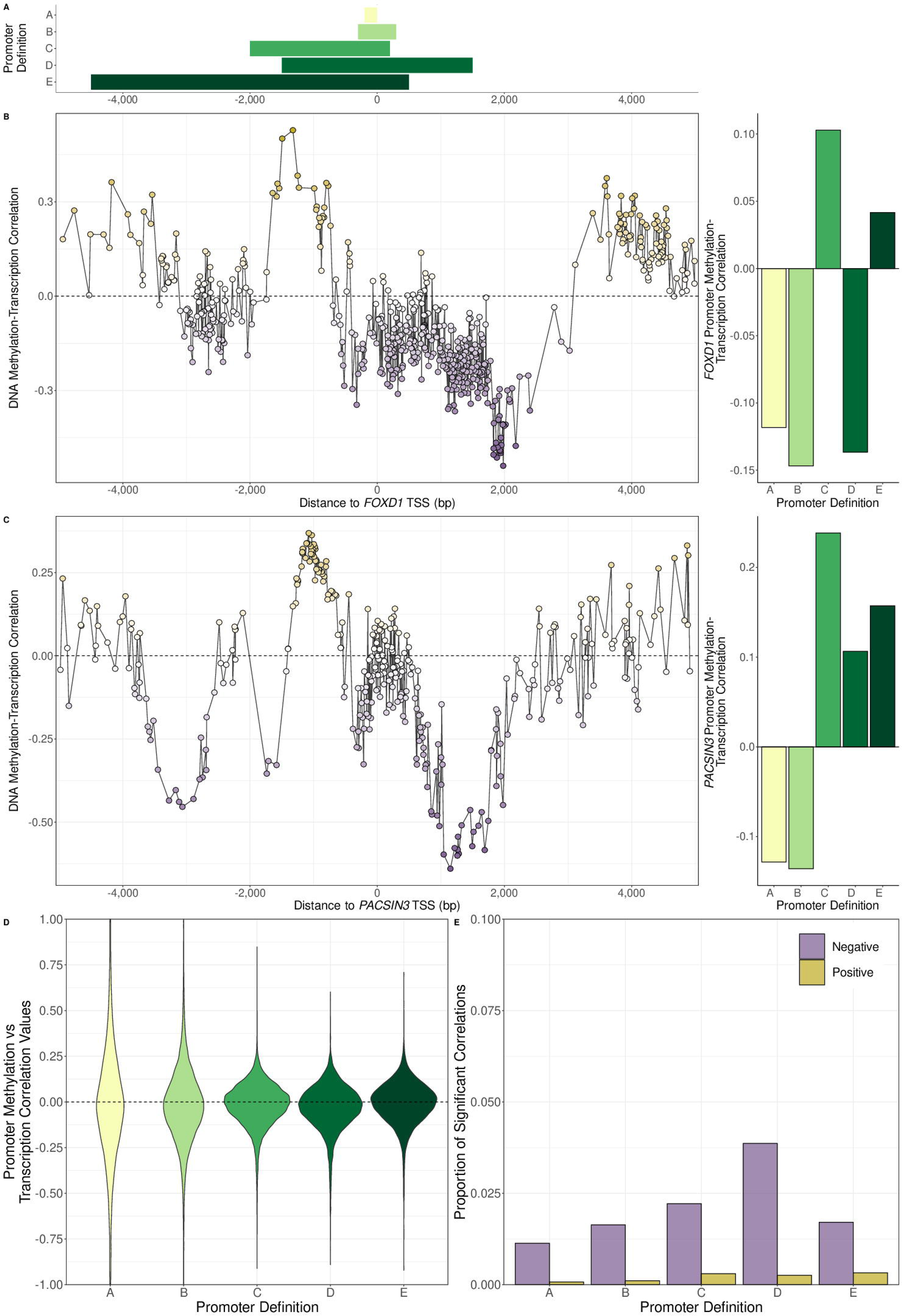
(A) The location of five different promoter definitions relative to the TSS from the same five different studies of DNA methylation as in Figure 1A. (B and C) The effect of the choice of promoter definition on promoter methylation-transcription correlations for *FOXD1* (using the TSS associated with the ENST00000615637 transcript) and *PACSIN3* (using the TSS associated with the ENST00000298838 transcript) in prostate metastasis samples. In the left-hand plots, the x-axes show distance of CpG sites upstream and downstream of the associated TSS in base pairs and the y-axes show Spearman correlation values between CpG methylation and transcriptional activity of the TSS. The right-hand plots show the promoter DNA methylation-transcription Spearman correlation values were calculated for the different promoter definitions using the average methylation values across CpG sites overlapping each definition. Different promoter definitions can result in opposing promoter methylation-transcription correlation values, though no correlation values were statistically significant. (D) The distribution of promoter methylation-transcription Spearman correlation values for all protein-coding transcripts using each of the 5 different promoter definitions in normal prostate samples. Most correlation values are close to zero. (E) The proportion of statistically significant correlations in normal prostate samples for each promoter definition divided into negative and positive correlations. Only a small minority of correlations are significant using any promoter definition. Among the significant correlations are a small number of positive ones.

Nevertheless, we wanted to determine if one definition generally tended to result in the strongest correlations between promoter DNA methylation and transcription. Thus, we calculated these correlations for protein-coding transcripts using each of the five chosen different definitions in the normal prostate samples (Figure 2D and E), prostate tumour and metastasis samples (Supplementary Figure 9). We found that, regardless of the promoter definition used, the correlation values were generally quite low, with about half of all correlation values having an absolute value less than 0.1 for all definitions across all datasets (Figure 2D and Supplementary Figure 9A and 9B) and only a very small proportion of all correlation values being statistically significant (Figure 2E and Supplementary Figure 9C and 9D). Promoter definition D, the definition with the greatest amount of sequence downstream of the TSS, resulted in the greatest number of significant correlation values in all groups of samples. This suggests that in addition to upstream regions, the region immediately downstream of the TSS could also be important to controlling transcriptional regulation via DNA methylation. Additionally, we observed that a small number of the statistically significant correlations are actually positive, supporting previous reports of positive association between DNA methylation and gene expression in certain settings (9, 10). Additionally, there were around twice as many significant correlation values overall in the prostate tumour samples and prostate metastases compared to normal prostate samples, possibly reflecting transcriptional dysregulation associated with aberrant DNA methylation in prostate cancer.

### Identification of transcript-proximal methylation-associated regulatory sites

Due to the frequent failure of fixed promoter definitions to capture the regions where DNA methylation is most strongly correlated with transcription, we sought to develop an alternative approach that would systematically identify these regions. Thus, we developed Methodical, a computational approach that uses WGBS and RNA-seq to identify such regions (Figure 3; details in Materials and Methods). We refer to these regions as transcript-proximal methylation-associated regulatory sites (TMRs). TMRs are classed as either having a negative or positive direction based on the sign of the correlation values between methylation of their CpG sites and transcription. Figure 4A and 4B show TMRs identified for *FOXD1* and *PACSIN3* in prostate metastasis samples.

**Figure 3:**
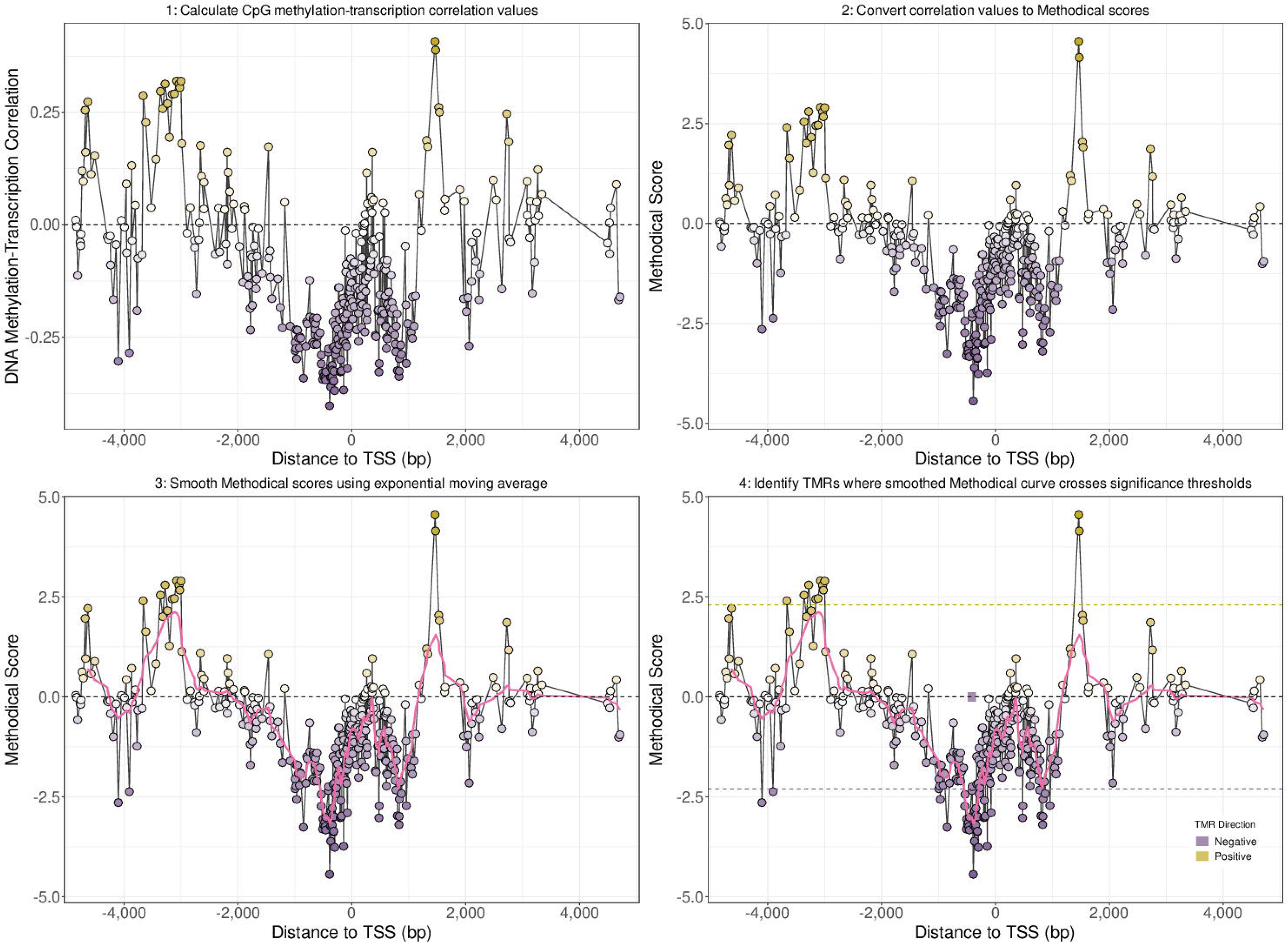
Overview of the identification of TMRs by Methodical using the TUBB6 transcript ENST00000591909 as an example. X-axes show distance from the ENST00000591909 TSS in base pairs. Y-axis of the top left panel indicates the Spearman correlation between DNA methylation levels of CpG sites and expression of ENST00000591909, while Y-axes for other panel indicates Methodical scores associated with these correlations. 1: First Spearman correlation values are calculated between methylation of CpG sites close to a TSS and the expression of the transcript associated with that TSS. 2: Next, these correlation values are converted into Methodical scores by taking the logarithm base 10 of the p-values associated with the correlation values multiplied by -1 if the correlation is positive. 3: These Methodical scores are then smoothed by using an exponential moving average, with the smoothed scores indicated by the pink line. 4: Finally, two significance thresholds are used to identify TMRs, one for positive TMRs indicated by the dashed gold line and another for negative TMRs indicated by the dashed purple line. The location of the identified negative and positive TMRs are shown by the purple and gold blocks, respectively.

**Figure 4:**
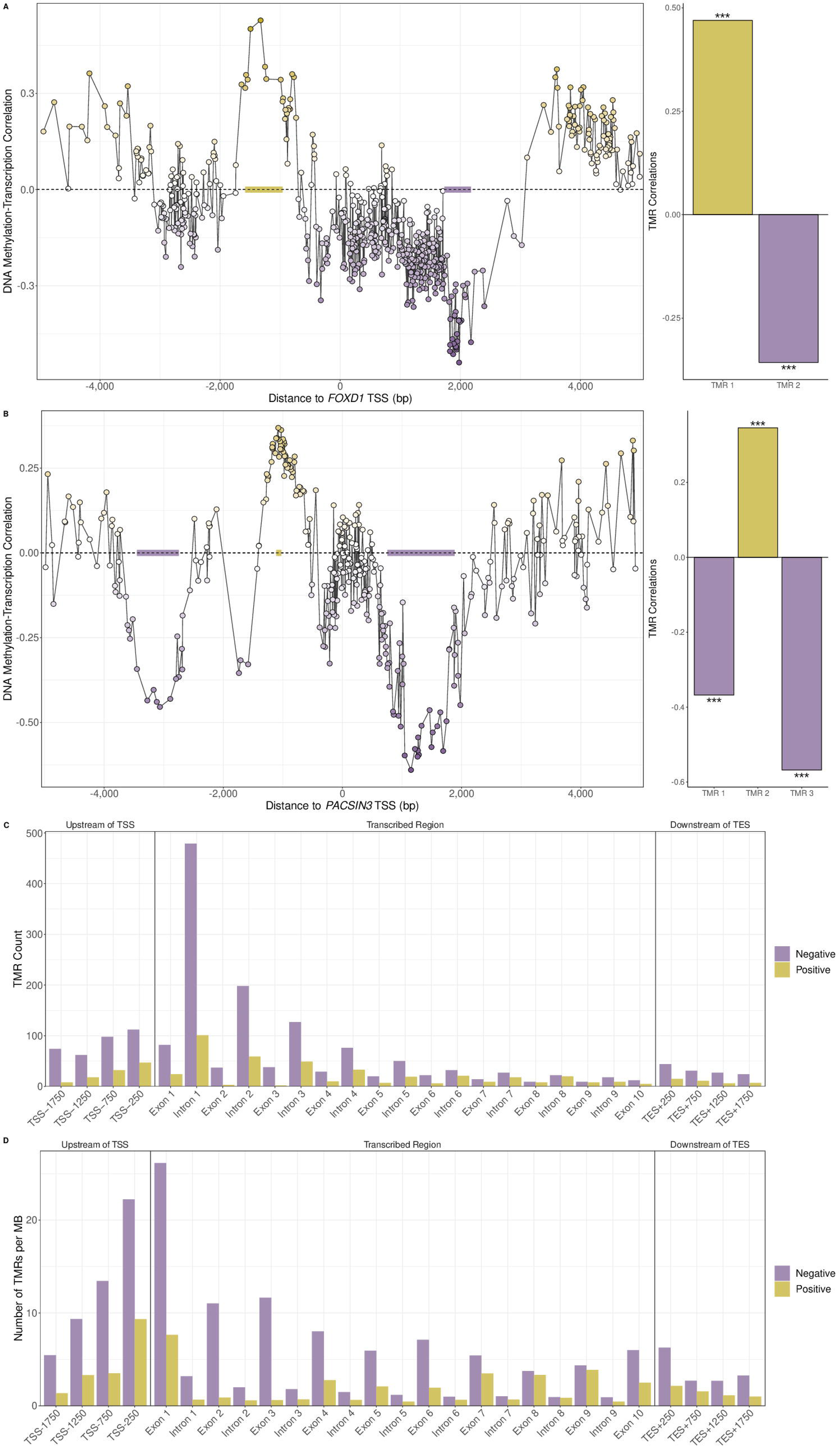
(A and B) The location of TMRs for *FOXD1* (associated with the ENST00000615637 transcript) and *PACSIN3* (using the ENST00000298838 transcript) found in prostate metastasis samples. X-axes in left-hand plots show distance of CpG sites from TSS in base pairs while Y-axes show Spearman correlation values between CpG methylation levels and transcription. The right-hand plots show the TMR DNA methylation-transcription Spearman correlation values calculated by averaging the methylation values across CpG sites overlapping each TMR. Stars indicate the level of statistical significance of correlations using a t-distribution. *FOXD1* has one negative and one positive TMR and *PACSIN3* has two negative and one positive TMR, all of which are significantly correlated with transcriptional activity. (C) Location of TMRs found in normal prostate samples within transcribed regions or within 5 kb upstream of the TSS or 5 kb downstream of the TES for all transcripts with 10 or fewer exons. Transcribed regions were separated into individual introns and exons while upstream and downstream regions were divided into 500 bp bins with the x-axis label indicating the centre of the bins. (D) The number of TMRs normalized by the number of MB covered in the genome by the indicated regions. Depending on whether the number of TMRs is normalized or not, either the first exon or first intron is the most common location for TMRs.

We first searched for TMRs within +/- 5 kilobases (kb) of TSS to gauge the typical distance of TMRs from the TSS and noted that most TMRs were found in the transcribed region just downstream of the TSS (Supplementary Figure 10A). We then decided to expand the search region for TMRs to the whole transcribed region for each transcript in addition to 5 kb upstream of the TSS and 5 kb downstream of the transcription end site (TES) in order to comprehensively study TMRs within transcribed regions and the regions upstream and downstream. We found 2, 022 TMRs in normal prostate samples, 2, 794 in tumour samples and 1, 623 in prostate metastases, about 25-30% of which were positive depending on the dataset. These were associated respectively with 1, 254, 1, 808 and 1, 146 transcripts and 1, 140, 1, 625 and 1, 086 genes (Supplementary Figure 10B).

We then investigated the distribution of TMRs among different regions overlapping and near to transcribed regions, including regions upstream of the TSS, introns, exons and regions downstream of the TES, We noted that just downstream of the TSS was the most common location for TMRs. Depending on whether we normalized classes of transcript-associated genomic regions (e.g. first exon, first intron, etc.) by the number of MB covered by the regions or not, either the first exon or first intron displayed the highest number of TMRs. Figure 4C shows the total number of TMRs found in different classes of transcript-associated genomic regions in normal prostate samples, plotting only TMRs associated with transcripts with 10 or fewer exons for display purposes. The first intron is by far the most common location of TMRs. However, as introns are on average much longer than exons, with the first intron being particularly long in general, it could be expected that simply due to its size, more TMRs would be found within the first intron. We thus decided to examine the number of TMRs in different regions after normalizing by the number of MB in the genome covered by these regions. Figure 4D shows the normalized number of TMRs for these regions, with the first exon then becoming the most common location, followed by the region from 500 bp upstream to the TSS and TMRs becoming less frequent further away from the TSS.

Thus, the first intron is the most common location of TMRs in absolute terms though the first exon is the region with the highest density of TMRs. We saw very similar patterns in the prostate tumour and prostate metastasis samples (Supplementary Figure 11). We also observed a similar distribution pattern when we examined the spatial distribution of TMRs we identified in a small number of diverse tissue samples from the Roadmap Epigenomics project (Supplementary Figure 12), indicating that is a general pattern for TMRs across tissues.

The presence of several TSS in close proximity regulating different isoforms of the same gene could potentially complicate the interpretation of the spatial distribution patterns of TMRs. To determine if we saw the same general pattern when examining only one TSS per gene, we selected the TSS associated with the Matched Annotation from NCBI and EBI (MANE) transcripts, a set of ENSEMBL transcripts comprising a high confidence transcript for each gene with complete sequence identity with a Refseq transcript. When we examined the distributions of the subset of TMRs associated with these TSS, we observed almost identical distribution patterns to those when considering TMRs associated with all TSS (Supplementary figure 13).

We found that there was a much higher overlap between the TMRs found in prostate tumour and prostate metastasis samples than that between the TMRs found in either group with those found in normal prostate samples. This is despite the fact that the tumour and metastasis samples come from different patient cohorts, while the tumour samples and normal prostate samples are from the same individuals. The transcripts and genes associated with TMRs displayed a similar pattern (Supplementary Figure 10C). This indicates that the TMRs identified in prostate tumours and metastases may be associated with transcriptional networks important to the development and progression of prostate cancer. Indeed, when we tested if the genes associated with TMRs in prostate tumour or metastases but not in normal samples were overrepresented for any KEGG pathways, we found that the terms “pathways in cancer” and “basal cell carcinoma” were enriched, strongly supporting a role for these genes in cancer development (one-sided Fisher’s exact test FDR corrected p-value < 0.05).

We also noted that negative TMRs from one dataset generally had a higher overlap with the negative TMRs than with the positive TMRs from the other two datasets and vice versa for positive TMRs, supporting a consistent association between TMR methylation and transcription across datasets. To further investigate this, we then evaluated the TMR methylation-transcription correlations across the three datasets, comparing the correlations for TMRs identified within a particular dataset, which we refer to as internal TMRs for that dataset, with those identified in another dataset, which we refer to as external TMRs for that dataset. For example, the TMRs identified in normal prostate samples are internal to normal prostate samples and external to prostate tumours and metastases.

Unsurprisingly, the correlation values for internal TMRs were very strong and almost all statistically significant (Supplementary Figure 10D and E). However, we also saw that a large proportion of correlation values involving external TMRs were strong with 40% statistically significant overall, a far higher proportion than with any fixed promoter definition (Figure 2E). This proportion increased to 49% when considering the TMRs identified in prostate tumour samples in prostate metastasis samples or vice versa, further supporting that the TMRs identified in prostate tumours and metastases are regions important to prostate tumorigenesis.

### Overlap with TMRs with Regulatory Features, Chromatin States and Transcriptional Regulator Binding sites

It has been reported that the relationship between DNA methylation and transcription varies with genomic context (5, 45) and also has opposing effects on the binding of different transcription factors (46). We thus decided to investigate if TMRs were associated with certain regulatory elements, chromatin states and transcriptional regulator binding sites.

We calculated the normalized number of TMRs overlapping different classes of genomic regulatory elements. We found that CpG islands, CpG shores and predicted enhancer regions were the regulatory elements with the greatest enrichment of both negative and positive TMRs (Figure 5A).

**Figure 5:**
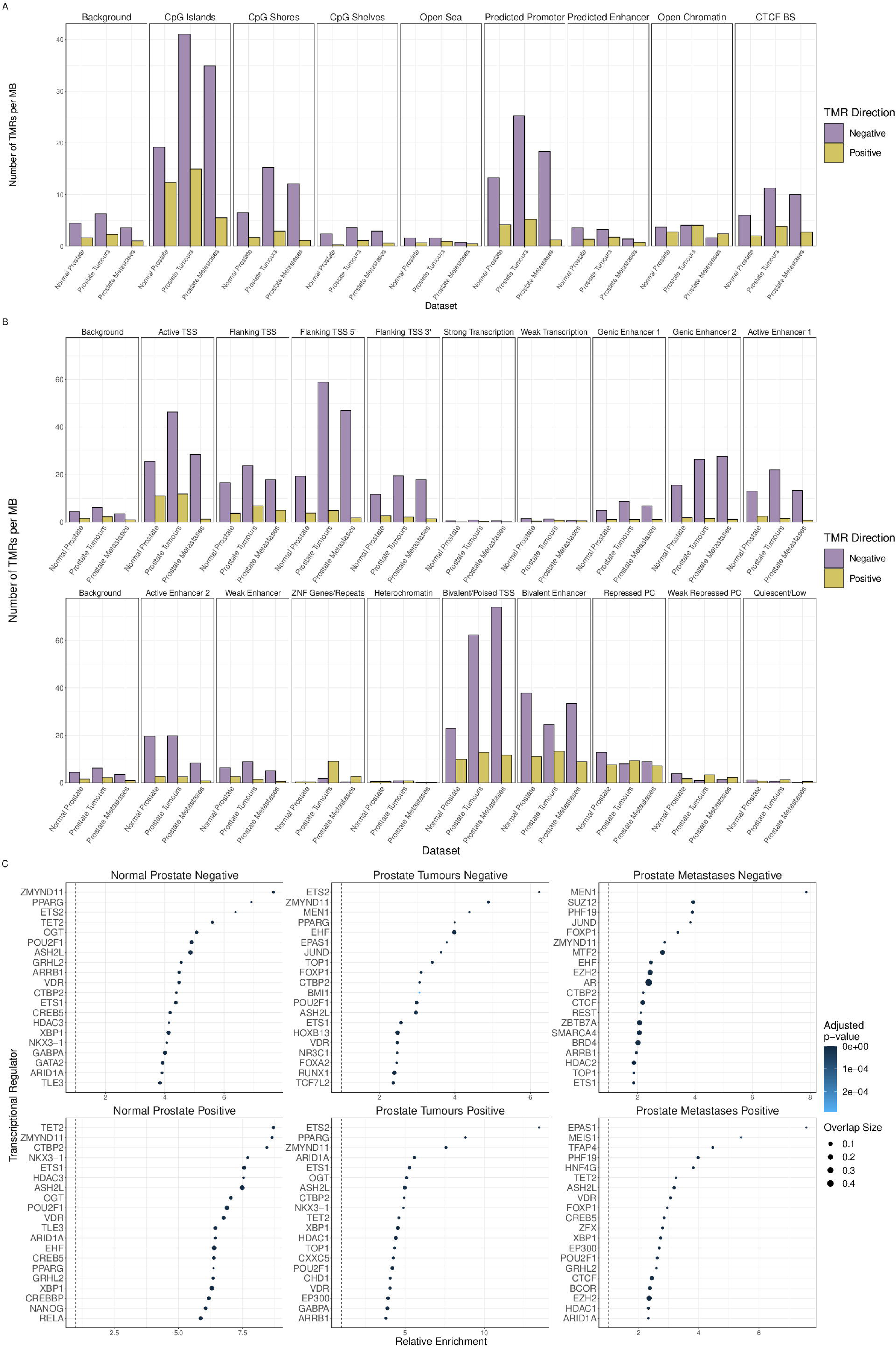
(A) The normalized number of TMRs overlapping different classes of genomic regulatory elements. Normalization was done to account for the fact that different classes of regulatory elements occupy varying proportions of the genome and was performed by dividing the number of TMRs overlapping each regulatory class by the number of MB of DNA in TMR search regions (the transcription units with 5 kb added upstream and downstream) overlapping the respective regulatory class. CTCF BS stands for CTCF binding site. (B) The normalized number of TMRs overlapping different chromatin states, calculated similarly as with the genomic regulatory elements above. (C) Enrichment of binding sites for transcriptional regulators among different groups of TMRs. The top 20 most enriched regulators for each TMR group is shown. Supplementary Table 1 lists all transcriptional regulators with significantly enriched binding sites.

Following on from this, we evaluated in a similar manner the overlap of TMRs with 18 chromatin states identified in healthy prostate tissue (47). The most common chromatin states associated with TMRs were the active TSS and flanking TSS 5’ states, followed by different enhancer states as well as bivalent states (bivalent/poised TSS and bivalent enhancer) (Figure 5B). Although we found that the most common location for TMRs was the region immediately downstream of the TSS, we found many more TMRs in the chromatin state labelled flanking TSS 5’ than in the state labelled flanking TSS 3’. We found that regions assigned to the flanking TSS 3’ state tended to be located much further from annotated TSS than regions assigned to the flanking TSS 5’ state and that neither state was preferentially located upstream or downstream of TSS, likely explaining this apparent contradiction.

We made similar observations when we examined the overlap of TMRs we identified in the samples from the Roadmap Epigenomics project with regulatory features and chromatin states, with TMRs enriched for CpG islands, predicted promoter regions and bivalent chromatin states (Supplementary Figure 14A and B).

Next we evaluated the overlaps of TMRs with binding sites in prostate tissue for 77 transcriptional regulators, including various transcription factors. The majority of these had significant overlaps with all categories of TMRs. MEN1, a known tumour suppressor gene associated with endocrine tumours (48), was one of the transcriptional regulators most strongly associated with negative TMRs, being the most enriched regulator for negative TMRs identified in prostate metastases and the third most enriched regulator for those identified in prostate tumours. Forkhead box (FOX) transcription factor family members FOXA2 and FOXP1 were also among the proteins with binding sites most strongly associated with negative TMRs and have both previously been implicated in prostate cancer (49, 50). Notably, binding sites for several chromatin remodelers were enriched for different TMR groups, including HDAC1, HDAC2, HDAC3, ARID1A, EZH2, SUZ12 and TET2 (Figure 5C; See Supplementary Table 1 for full results).

### Methylation Change at TMRs in Tumorigenesis

DNA Methylation change affects much of the genome in cancer, with increases of methylation at specific loci, particularly CpG islands, contrasting with a general loss of methylation across the rest of the genome (51–53). Given the vast number of regions affected by DNA methylation change, it has been difficult to determine which changes are playing an active role in tumour development and progression (54). Considering this, we reasoned that examination of methylation change at TMRs in cancer could help identify regions where altered methylation is associated with transcriptional changes in cancer and thus more likely to be clinically relevant.

We thus tested differential methylation of TMRs between the prostate tumour and matching normal prostate samples and discovered that TMRs are very frequently affected by altered methylation in prostate cancer (Figure 6A). Intriguingly, negative and positive TMRs identified in prostate tumours or metastases tended to display opposing methylation change in prostate tumours, with negative TMRs often becoming hypermethylated and positive tumours often becoming hypomethylated.

**Figure 6:**
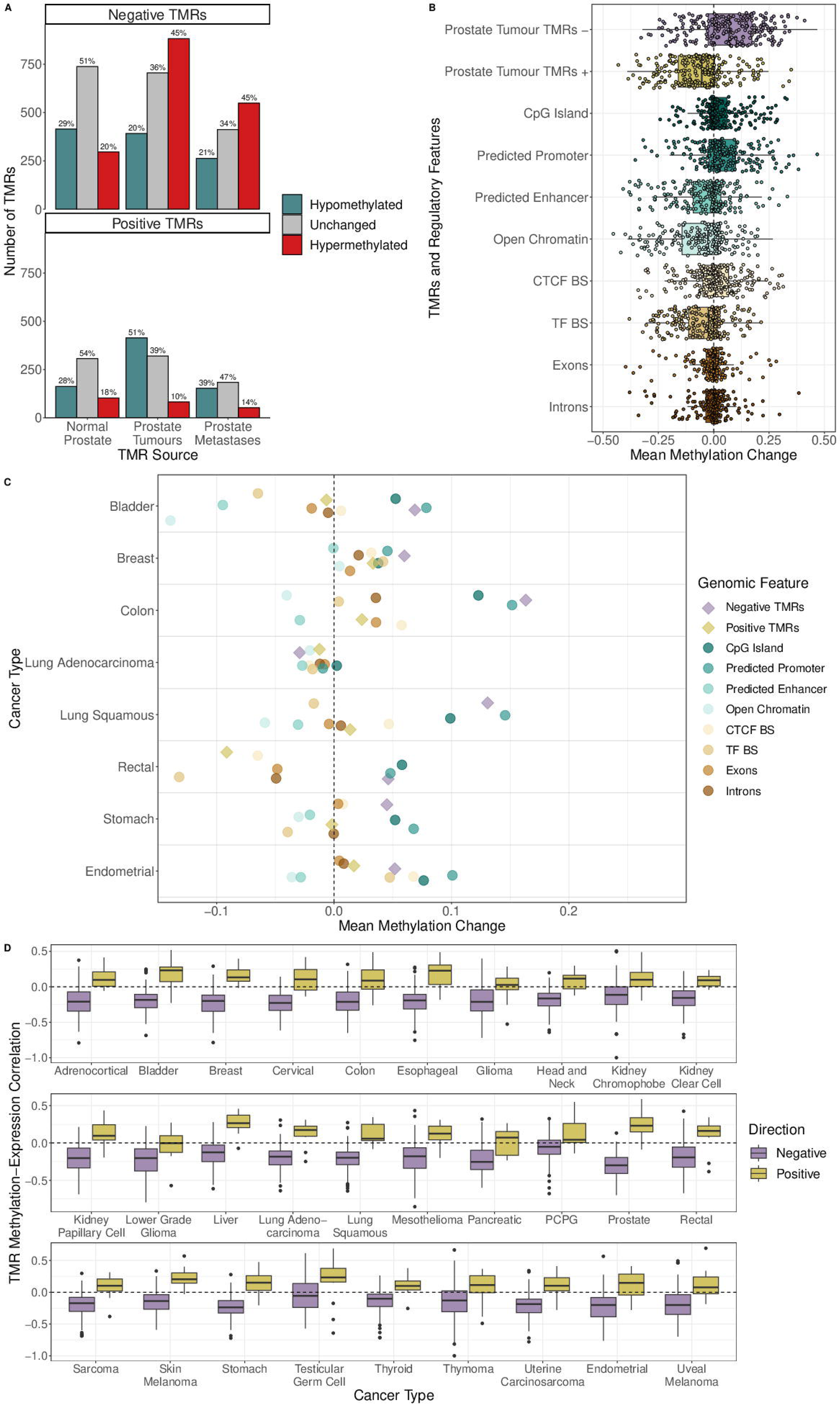
(A) The number of differentially methylated negative and positive TMRs in prostate cancer. The x-axis shows the source of TMRs, the y-axis the number of TMRs and colour indicates the direction of methylation change. Percentages above bars indicate the percentage of all TMRs in the relevant group affected by the indicated methylation change. (B) Methylation change at TMRs in prostate tumours compared to different regulatory regions. All classes of genomic regions were subset for regions not overlapping repeats. Boxplots show the distributions of the mean methylation change of regions belonging to the indicated class. The mean methylation change of 250 random regions belonging to each class are plotted as points on top of the boxplots. TMRs display a strong tendency to become hypermethylated, particularly negative TMRs identified in prostate tumour and metastases samples. (C) Methylation change at various classes of genomic regions across in eight cases of different cancer types with WGBS data from TCGA. TMRs are those identified in prostate metastasis samples and are indicated by diamonds with other classes of regions represented by circles. All classes of genomic regions were subset for regions not overlapping repeats. TMRs are the most hypermethylated regions in the breast cancer and colon cancer cases. (D) Distribution of Spearman correlation values between methylation of TMRs associated with MANE transcripts identified in prostate tumour samples and expression of the relevant gene across different cancer types from TCGA. Direction of correlation values are generally as would be expected, with negative correlations for negative TMRs and positive correlations for positive TMRs, though the strongest correlations were generally observed in prostate cancer samples from TCGA. PCPG stands for Pheochromocytoma and Paraganglioma.

Differentially methylated regions (DMRs) were previously reported in the prostate metastasis samples (29) and so we were interested in the overlap between these regions and the TMRs we identified in those samples. The vast majority (96%) of the DMRs were hypomethylated, in contrast to the subset of DMRs which overlapped negative TMRs, where 35% were hypermethylated. We hypothesized that the increased methylation at TMRs is possibly more likely to be functionally relevant than the loss of methylation at the majority of hypomethylated DMRs. Supporting this, when we performed gene set overrepresentation analysis for the nearest genes to all DMRs within our TMR search space (transcribed regions +/- 5 kb), we found no significantly enriched KEGG or MSigDB Hallmark pathways. However, when we tested the genes associated with subset of DMRs that overlapped TMRs, we found several enriched KEGG pathways, including “pathways in cancer”, “focal adhesion” and “ECM receptor interaction”, and also various hallmark pathways, including “androgen response”, “epithelial mesenchymal transition”, “KRAS signaling up” and “Notch signaling” (one-sided Fisher’s exact test, FDR-corrected p-value < 0.05).

We also wondered how often DMRs discovered near the vicinity of transcribed regions are associated with transcriptional activity. Thus, we selected for all DMRs which overlapped the regions in which we searched for TMRs (transcribed regions plus 5 kb upstream and downstream) and calculated the correlation between the methylation of these DMRs and expression of the associated transcript. We found that less than 10% of this subset of DMRs were significantly associated with expression. Thus, standard differential methylation testing can lead to the identification of a vast number of regions which are generally not correlated with the expression of nearby genes. Consequently, methylation change at TMRs may be more functionally relevant than at DMRs in general, supported by the overrepresentation of genes in cancer-related KEGG and Hallmark pathways among TMRs overlapping DMRs.

We next compared the methylation change at TMRs to that at different regulatory regions. Negative TMRs identified in prostate tumours were the most hypermethylated of all classes of genomic regions studied, displaying an even greater degree of methylation gain than CpG islands or predicted promoter regions (Figure 6B). Conversely, positive TMRs identified in prostate tumours were the class of region with the greatest loss of methylation.

We subsequently performed pathway overrepresentation analysis for the genes associated with differentially methylated TMRs using KEGG pathways. Strongly supporting a role for altered TMR methylation in tumorigenesis, we found that the term “pathways in cancer” was enriched among the genes associated with negative TMRs identified in prostate tumours and metastases which were hypermethylated in prostate cancer. (Supplementary Figure 15A).

Accordingly, we found several examples of TMRs associated with prostate cancer-related genes displaying methylation changes in prostate tumours. This included hypermethylation of negative TMRs associated with tumour suppressor genes, such as *BCOR and SOCS1*, metastasis suppressors such as *CD44* and *TIMP1*. Additionally, *GSTP1,* one of the most frequently hypermethylated genes in prostate cancer and a proposed caretaker gene (55), displayed pronounced hypermethylation of a TMR located just downstream of its TSS. Conversely, a positive TMR near the TSS of *TNFAIP8*, an androgen response gene playing an anti-apoptotic role in prostate cancer (56), also exhibited a spike of increased methylation. These TMRs varied in their location relative to the TSS, with some overlapping the TSS, others located upstream and others located downstream (Supplementary Figure 16).

Several other KEGG pathways related to the extracellular matrix, such as “focal adhesion”, “regulation of actin cytoskeleton” and “ECM receptor interaction” were also enriched, suggesting that differential methylation of TMRs could be associated with tumour invasiveness.

Conversely, genes associated with hypomethylated negative TMRs were enriched for the androgen response pathway and related pathways such as cholesterol homeostasis and estrogen response, indicating that loss of methylation at these TMRs is associated with increased androgen signalling (Supplementary Figure 15B). Only a few pathways were significantly enriched among genes associated with positive TMRs, probably related to the identification of much fewer positive TMRs than negative TMRs, but included terms related to angiogenesis KRAS signalling and also EMT.

To investigate if methylation change at TMRs may be a common occurrence across different cancers, we evaluated methylation change at the TMRs identified in the prostate metastasis samples in tumour and matching normal samples with WGBS data from eight different patients with different cancer types from TCGA (57). We found that the negative TMRs once again often gained methylation, being the class of region with the greatest methylation increase in the breast cancer and colon cancer cases (Figure 6C). Finally, we wanted to confirm if the methylation of the TMRs we identified displayed the expected associations with gene expression in different cancer types from TCGA. However, only gene-level expression data, but not transcript-level, is available publicly from TCGA, though TMRs are specifically associated with the expression of individual transcripts. Thus, we decided to evaluate only TMRs associated with MANE transcripts. The MANE transcripts are generally the most expressed transcripts for each gene, and so we reasoned that the the methylation of TMRs associated with MANE transcripts should generally also be strongly associated with the overall expression for the corresponding gene.

Additionally, the vast majority of the DNA methylation data for TCGA samples was produced using the Illumina 450K methylation array which covers only a minority of CpG sites in the genome. Thus, we evaluated the correlation values between the methylation of TMRs associated with MANE transcripts identified in prostate metastasis samples which also overlapped CpG sites covered by the Illumina 450K methylation array and expression of the relevant gene in 29 different cancer types from TCGA. Only 656 TMRs (571 negative and 85 positive) were associated with MANE transcripts and were targeted by one or more probes from the array. Additionally, methylation data was generally only available for a proportion of the CpG sites in each of these TMRs. Regardless, we observed that the signs of the correlation values generally matched those expected given the direction of TMRs; negative TMRs generally displaying negative correlations and positive TMRs generally exhibiting positive correlations (Figure 6D). Thus, many of the TMRs we identified in prostate cancer are regions where methylation is associated with gene expression in a pan-cancer manner. We did observe that the strongest correlations were in the prostate cancer samples from TCGA, indicating that there is some tissue-specificity to the associations. Therefore, applying Methodical to datasets with WGBS and RNA-seq for other tissues could lead to the identification of TMRs which are more relevant to different tissue or cancer types.

### Methylation Microarrays Often Miss Regions Where DNA Methylation Is Most Strongly Associated with Transcription

Several previous studies of the association between DNA methylation and transcription have used methylation data produced by Illumina arrays (10, 27, 28) and so we wanted to evaluate if analysis of the CpG sites targeted by these arrays could adequately capture the general correlation patterns between DNA methylation and transcription around TSS. We first noted that the CpG sites covered by Illumina arrays were biased towards those in within about 1 kb upstream of the TSS, with relatively low coverage of CpGs downstream of the TSS where we identified the greatest number of TMRs, (Figure 7A and B).

**Figure 7:**
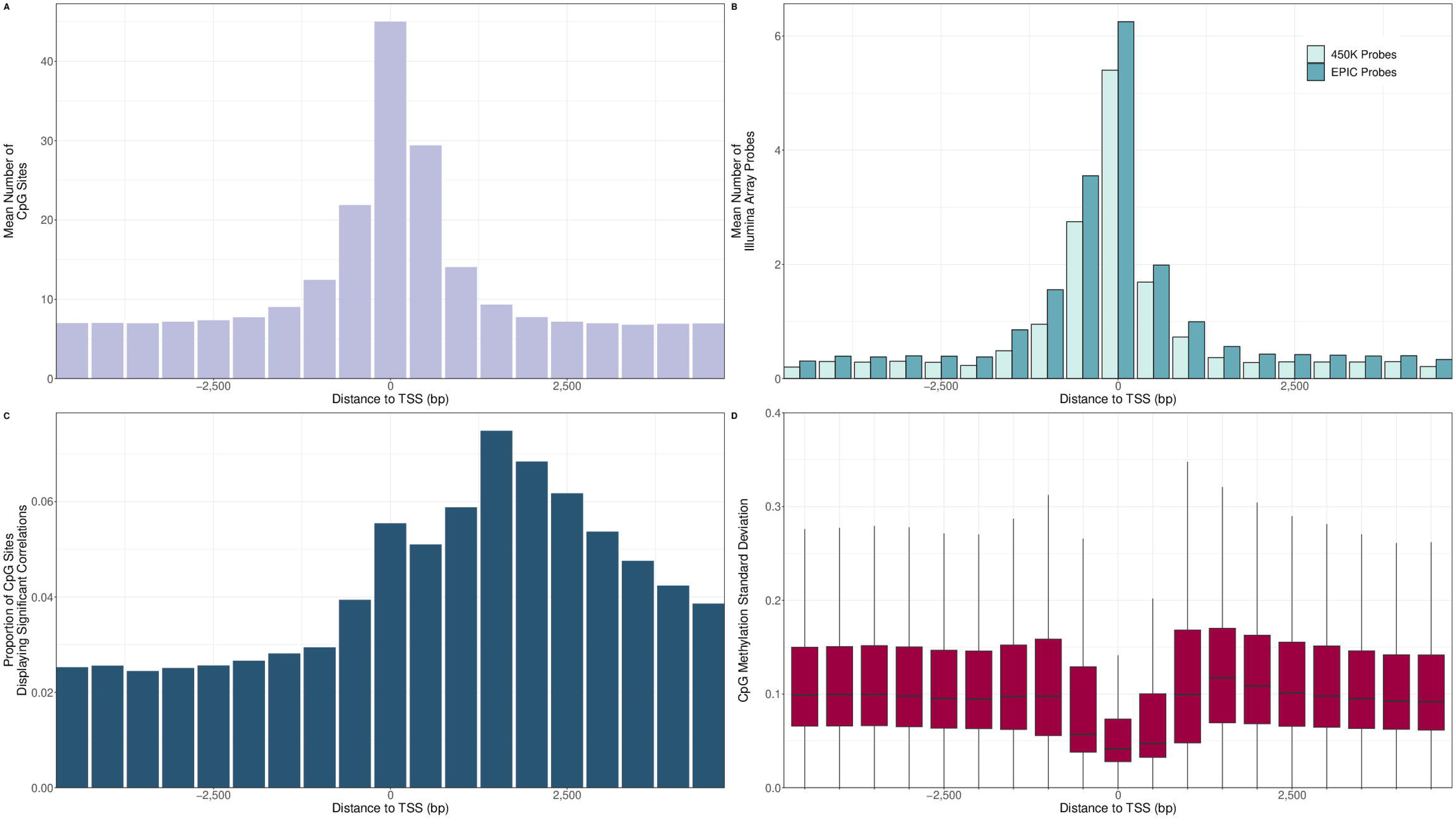
(A) The mean number of CpG sites in 500 bp bins around TSS. The bin overlapping the TSS is the most common location for CpG sites, with the bin just downstream the second most common. (B) The mean number of CpG sites targeted by the Illumina HumanMethylation450 and EPIC methylation microarrays in 500 bp bins around TSS. Most CpGs targeted are in the bin overlapping the TSS or the bin 500 bp upstream, with much fewer CpGs targeted elsewhere, including the region just downstream of the TSS with a high concentration of CpG sites. (C) The proportion of CpG sites where DNA methylation was significantly correlated with transcript expression in prostate tumour samples in 500 bp bins around the TSS. Significant was defined as a corrected p-value less than 0.05 for the Spearman correlation values. The greatest proportion of significant correlation is found downstream of the TSS. (D) Standard deviation values for the methylation of CpG sites in 500 bp bins around TSS. CpGs in the region around the TSS generally have the lowest standard deviations, while those located between 1-2 kb downstream tend to have the highest standard deviations.

When we then examined the distribution of CpGs where DNA methylation was significantly correlated with transcription in prostate tumour samples, we found that the region with the greatest proportion of CpGs with significant correlations was the region downstream of the TSS (Figure 7C), fitting with this being the region where we discovered the most TMRs. Relatedly, when we examined the standard deviation of methylation of CpG sites grouped by distance to TSS, we found that those CpG sites downstream of the TSS tended to have higher standard deviations than the regions upstream (Figure 7D). We made similar observations with WGBS data in normal prostate samples (Supplementary Figure 17A and B) and prostate metastasis (Supplementary Figure 17C and D), though with the highest proportion of significant correlations in normal prostate samples found at the TSS. We also noted a similar pattern in normal and tumour samples using Illumina methylation array data from several different cancer types from TCGA (Supplementary Figure 18), demonstrating this is a general pattern across tissue and cancer types.

Thus, the bias of methylation arrays to the region immediately surrounding the TSS and just upstream has likely resulted in an underappreciation of the association of DNA methylation further from the TSS with transcription, particularly downstream methylation. Indeed, we found that, depending on the dataset, 50-70% of TMRs we identified did not contain a single CpG site that was targeted by the Illumina HumanMethylation450 array.

## Discussion

Here, we have demonstrated firstly how the use of different arbitrary promoter definitions in studies of DNA methylation can lead to profoundly discordant results. These different promoter definitions can result in vastly different numbers of differentially methylated promoters being identified and perhaps even more strikingly, can lead to completely opposing results even at the same TSS. We found that these contradictory results with different promoter definitions can affect thousands of different TSS. In light of this, the wide variation in how promoters are defined between different studies of DNA methylation is concerning.

Unsurprisingly given the effect of promoter definition choice on the calculation of promoter methylation levels, we also found that the correlation values between promoter methylation levels and transcript expression are hugely influenced by the choice of promoter definition. The vast majority of these correlation values are very weak and statistically insignificant, consistent with other studies modelling gene expression using DNA methylation (7, 58–61), and we have revealed that this is in part because fixed-size promoter definitions often miss the complex relationship between methylation of CpG sites, distance from TSS and transcriptional activity. Regions where DNA methylation is negatively correlated, positively correlated or uncorrelated with transcription can all occur in the region immediately upstream and downstream of the TSS, with different TSS displaying completely different patterns.

This is in line with previous work extracting higher order features from methylation data, which enabled the identification of different five major spatially correlated promoter methylation patterns (14). Unsurprisingly, a uniformly hypermethylated pattern was associated with transcriptional repression. However, a uniformly lowly-methylated pattern was also unexpectedly associated with repression. Conversely, a U-shaped pattern characterized by a central hypomethylated region flanked by hypermethylated regions on either side was associated with high expression and interestingly this pattern was largely found at CpG islands and often associated with housekeeping genes. This pattern supports the idea that hypermethylation in CpG shores is positively associated with transcription. A forward S-shaped pattern characterized by lowly methylated region which gradually transitions into a highly methylated region and the reverse S-shaped pattern were associated with an intermediate level of gene expression. Taken together, these patterns underscore that the effect of DNA methylation on transcriptional activity is highly context dependent.

Approaches using more sophisticated machine learning algorithms, such as support vector machines or deep learning approaches, with higher resolution methylation features have proven to predict transcriptional activity much more accurately than simpler linear approaches (14, 17, 18). The superiority of these complex models likely reflects the intricate relationship between methylation at different CpG sites in both promoter-proximal and distal enhancer regions and transcriptional activity. However, difficulty in interpreting such models makes determining the precise regions where DNA methylation is the most important for predicting transcriptional activity challenging. In contrast, our study specifically focuses on identifying and characterizing regions with the strongest correlations between DNA methylation and transcription.

In response to these issues, we believed that WGBS and RNA-seq data could be combined to enable the systematic identification of precise regions where DNA methylation is associated with transcription, providing an alternative to the use of arbitrary fixed-size promoter definitions. Hence we developed Methodical, a novel algorithm that integrates WGBS and RNA-seq data to identify regions where CpG methylation displays consistent correlation with transcriptional activity of associated TSS. We call these regions transcript-proximal methylation-associated regulatory sites (TMRs), with TMRs having either a negative or positive direction depending on the correlation between DNA methylation and transcription. We applied Methodical to three datasets, one for normal prostate, one for prostate tumours and another for prostate metastases, with a large number of samples with both WGBS and RNA-seq data to identify a set of TMRs for each dataset.

As should be expected, the methylation of TMRs displays a much stronger correlation with transcriptional activity than the methylation of fixed promoter definitions when evaluating the correlations within the dataset in which the TMRs were identified. However, the association of TMR methylation with transcriptional expression could often be validated in other datasets, including those for other tissue types. We did note that the TMRs we identified in prostate tumours displayed stronger correlations when evaluated in prostate cancer than in other cancer types from TCGA, indicating that TMRs are partially tissue specific.

We discovered that the most common location for TMRs was the region immediately downstream of the TSS. Supporting this, we also found that the region within the first 2 KB downstream of the TSS was generally the region with the greatest proportion of CpG sites where DNA methylation was significantly correlated with transcription. We observed this pattern in diverse tissue and cancer types using both WGBS and methylation array data. This is in line with other recent studies modelling transcription using DNA methylation which found that methylation of the region downstream of the TSS was the most important for predicting gene expression (15, 16).

∼25-30% of TMRs were positive, depending on the dataset, demonstrating that the conventional wisdom that DNA methylation is negatively associated with transcription does not always hold true and supporting previous reports of positive associations between DNA methylation and transcription (9, 10). Several possible mechanisms for transcriptional activation by DNA methylation have been proposed, including blocking the binding of transcriptional repressors and activation of alternative promoters (9).

We found that TMRs are enriched for certain genomic elements and chromatin states inferred from healthy prostate samples (47). Notably, negative TMRs from prostate tumours and metastases were highly enriched for a bivalent/poised TSS chromatin state. This bivalent state is characterized by the co-occurrence of both activating and repressive histone modifications, such as H3K4me3 and H3K27me3, respectively, and was first identified at the promoters of developmentally regulated genes in embryonic stem cells (ESCs). This led to the hypothesis that bivalency poises these developmental genes for rapid activation during development while maintaining a transcriptionally inactive state in the absence of activating signals (62, 63).

However this hypothesis has been challenged by the work of Kumar *et al.*(*64*). They demonstrated that bivalency does not seem to poise genes for rapid activation as the activation of bivalent genes was neither stronger nor more rapid than that of transcriptionally silenced genes lacking H3K4me3 at their promoters. Instead, they proposed that H3K4m3e may represent a general mechanism to maintain the unmethylated state of CGIs, even those associated with transcriptionally inactive genes.

While bivalent promoters generally display low methylation levels in normal cells (65, 66), it has been repeatedly observed that most genes that become hypermethylated in cancer have CGI promoters which are bivalently marked in ESCs (67–70). This suggests that bivalency predisposes these genes for methylation gain in cancer. Additionally, bivalent regions often display the strongest hypermethylation of all chromatin states (70, 71). However, as loss of bivalent chromatin seems to be a general feature of cancer (70), it may be that it is actually the loss of of bivalency(70), and H3K4me3 in particular, that leads to increased methylation, rather than the presence of bivalency *per se* (*64*). This was supported by the finding that regions that lost bivalency in cancer cell lines displayed greater hypermethylation than regions which remained bivalent (70). Thus, the prevalence of TMRs in bivalent regions may indicates that the gain in methylation in these regions is often associated with transcriptional repression of the associated genes.

Given the vast numbers of regions affected by DNA methylation change in cancer, we thought that focusing on methylation change at TMRs in cancer could improve our understanding of how DNA methylation alterations lead to transcriptional misregulation during tumorigenesis. We discovered that TMRs displayed a pronounced tendency to become hypermethylated in prostate cancer, though a substantial minority of them also became hypomethylated. Negative TMRs were affected by greater methylation change on average than all other classes of genomic elements studied, including CpG islands and predicted promoter regions. TMRs were also the most hypermethylated elements in some cases when we examined their methylation in other tumour types. This is fitting with a prior study reporting that regions displaying variable methylation between different tissue types and during reprogramming of induced pluripotent stem cells, and thus presumably associated with developmental gene expression networks, are often differentially methylated in a variety of cancer types (58).

Pathway overrepresentation analysis for genes associated with hypermethylated negative TMRs in prostate cancer revealed many pathways related to cancer, invasiveness or oncogenic signalling pathways, supporting a role for TMR hypermethylation in tumorigenesis. Conversely, hypomethylated negative TMRs were enriched for genes involved in the androgen response, fitting with previous observations of hypomethylation at genes involved in the androgen-response in prostate cancer (21, 29).

To our knowledge, this is one of the most detailed studies to date of the relationship between DNA methylation around TSS and transcriptional activity. Several other studies of the association between DNA methylation and transcription have used methylation data coming from Illumina methylation microarrays (10, 27, 28). These microarrays profile only a small fraction of CpG sites in the human genome and are highly biased towards certain genomic contexts, particularly upstream of the TSS. We have shown that regions downstream of the TSS actually have the greatest proportion of CpG sites where DNA methylation is significantly correlated with transcription. Thus, studies based on these arrays have missed many regions where DNA methylation is associated with transcription and likely have led to an underappreciation of the association of DNA methylation downstream of TSS with transcriptional activity. Future studies of DNA methylation should therefore aim for greater coverage of regions downstream of TSS.

Application of Methodical to other datasets with large numbers of samples with base-resolution methylation data and RNA-seq data for different tissue or tumour types when they become available would enable the identification of TMRs which may be more relevant for different tissue and cancer types. Here we only examined regions proximal to TSS, however Methodical could also be used to identify distal regions, such as enhancers, where DNA methylation is associated with transcription. In that case, care should be taken to account for correlations arising from gene co-expression, such as through a permutation-based approach (72). Furthermore, as Methodical can only identify correlative relationships, experimental studies would be needed to demonstrate that TMR methylation is causally involved in influencing transcription. For example, investigating if treatment with a demethylating agent such as 5-azacytidine or experimental alteration of TMR methylation using CRISPR-Cas9 technology leads to the expected expression changes could confirm if TMR methylation is causally involved in gene regulation.

In summary, Methodical enables the identification of regions where DNA methylation is correlated with transcriptional activity, enabling insights into the fundamental relationship between DNA methylation and gene expression. Better understanding of this relationship will in turn improve our understanding of how altered DNA methylation in cancer is associated with perturbation of the transcriptome during tumour development.

## Acknowledgements

The work leading to this manuscript was supported by Fondazione AIRC, grant reference numbers MFAG n. 21791 and Bridge Grant n. 29162 and Investigator Grant 30887. It has been partially supported by the Italian Ministry of Health with Ricerca Corrente and 5 × 1000 funds. The graphical abstract was created using BioRender.com.

We would also like to thank Dr. David Quigley and Dr. Jing Li for providing much of the data used in this study.

## Author contributions

R.H. performed the statistical and computational work. R.H. and M.S. conceived the study and wrote the manuscript. M.S. supervised the work.

## Data availiability

Processed WGBS data for normal prostate and prostate tumour samples from the CPGEA project ^12^ was downloaded from download.big.ac.cn:/gsa-human/HRA000099/processed/all_processed_files.tar.gz while processed WGBS data for prostate metastases from the MCRPC project (29) was provided by the authors. metastasis samples bedGraph files for WGBS analysis of 39 tumours and 8 matching normal samples from TCGA were downloaded from https://urlsand.esvalabs.com/?u=https%3A%2F%2Fzwdzwd.s3.amazonaws.com%2Fdirectory_listing%2FtrackHubs_TCGA_WGBS_hg38.html%3Fprefix%3DtrackHubs%2FTCGA_WGBS%2Fhg38%2Fbed%2F&e=8c00e339&h=dd5873df&f=y&p=n. Illumina HumanMethylation450 files with probe beta values for tumour and normal samples for cancer types from TCGA were downloaded from the Genomic Data Commons (GDC) using the GDC-client tool. Genomic locations of probes for Illumina methylation microarrays were downloaded from the Illumina website. BED files for WGBS data for 38 different non-cancer human tissue samples belonging to 19 different tissue types from the Roadmap Epigenomics project (65) were downloaded from the ENCODE data portal (73). See Supplementary Table 2 for files and tissue types. CpG methylation values and RNA-seq transcript counts for the CPGEA and MCRPC projects are available as a Bioconductor ExperimentHub package at https://bioconductor.org/packages/release/data/experiment/html/TumourMethData.html.

FASTQ files for RNA-Seq data from the CPGEA were downloaded from the Genome Sequence Archive for Human (http://bigd.big.ac.cn/gsa-human/). BAM files for RNA-seq data for the MCRPC project were downloaded from GDC using the GDC-client (https://portal.gdc.cancer.gov/repository) and FASTQ files were generated from them using Samtools (version 1.11). STAR count files with gene expression counts for TCGA were downloaded from GDC using the GDC-client for each cancer type. RNA-seq gene counts for Roadmap Epigenomics project samples were downloaded from the ENCODE data portal. See Supplementary Table 2 for files and tissue types.

Methodical is available as an R/Bioconductor package at https://bioconductor.org/packages/methodical

Supplementary Table 1 lists all significantly enriched transcriptional regulators for TMR groups. Supplementary Table 2 gives the tissue types and file accession ID for all samples from the Roadmap Epigenomics project used in the study.

BED files with coordinates of TMRs for the hg38 genome build are available to download from https://github.com/richardheery/heery_2024_scripts/blob/master/finding_tmrs/tmr_bed_files

Scripts to reproduce figures are available at https://github.com/richardheery/heery_2024_scripts

