## Supplementary Figures for "Systematic identification of regions where DNA methylation is correlated with transcription refines regulatory logic in normal and tumour tissues"

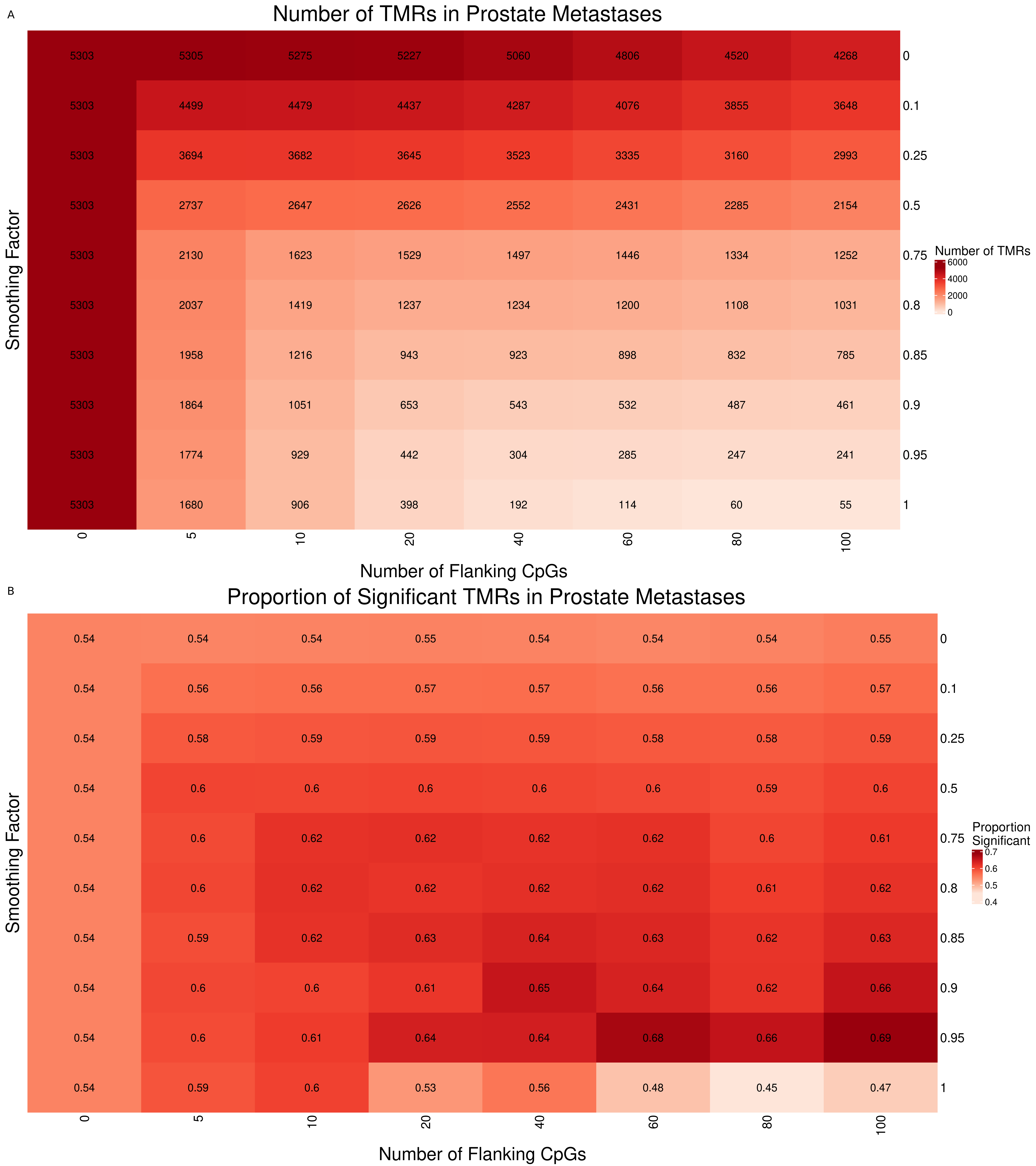
A
Number of TMRs in Prostate Metastases
5303 5305 5275 5227 5060 4806 4520 4268 0
5303 4499 4479 4437 4287 4076 3855 3648 0.1
5303 3694 3682 3645 3523 3335 3160 2993 0.25
5303 2737 2647 2626 2552 2431 2285 2154 0.5
5303 2130 1623 1529 1497 1446 1334 1252 0.75
5303 2037 1419 1237 1234 1200 1108 1031 0.8
5303 1958 1216 943 923 898 832 785 0.85
5303 1864 1051 653 543 532 487 461 0.9
5303 1774 929 442 304 285 247 241 0.95
5303 1680 906 398 192 114 60 55 1
0 5 10 20 40 60 80 100
Number of Flanking CpGs
Smoothing Factor
Number of TMRs
6000
4000
2000
0
B
Proportion of Significant TMRs in Prostate Metastases
0.54 0.54 0.54 0.55 0.54 0.54 0.54 0.55 0
0.54 0.56 0.56 0.57 0.57 0.56 0.56 0.57 0.1
0.54 0.58 0.59 0.59 0.59 0.58 0.58 0.59 0.25
0.54 0.6 0.6 0.6 0.6 0.6 0.59 0.6 0.5
0.54 0.6 0.62 0.62 0.62 0.62 0.6 0.61 0.75
0.54 0.6 0.62 0.62 0.62 0.62 0.61 0.62 0.8
0.54 0.59 0.62 0.63 0.64 0.63 0.62 0.63 0.85
0.54 0.6 0.6 0.61 0.65 0.64 0.62 0.66 0.9
0.54 0.6 0.61 0.64 0.64 0.68 0.66 0.69 0.95
0.54 0.59 0.6 0.53 0.56 0.48 0.45 0.47 1
0 5 10 20 40 60 80 100
Number of Flanking CpGs
Smoothing Factor
Proportion Significant
0.7
0.6
0.5
0.4

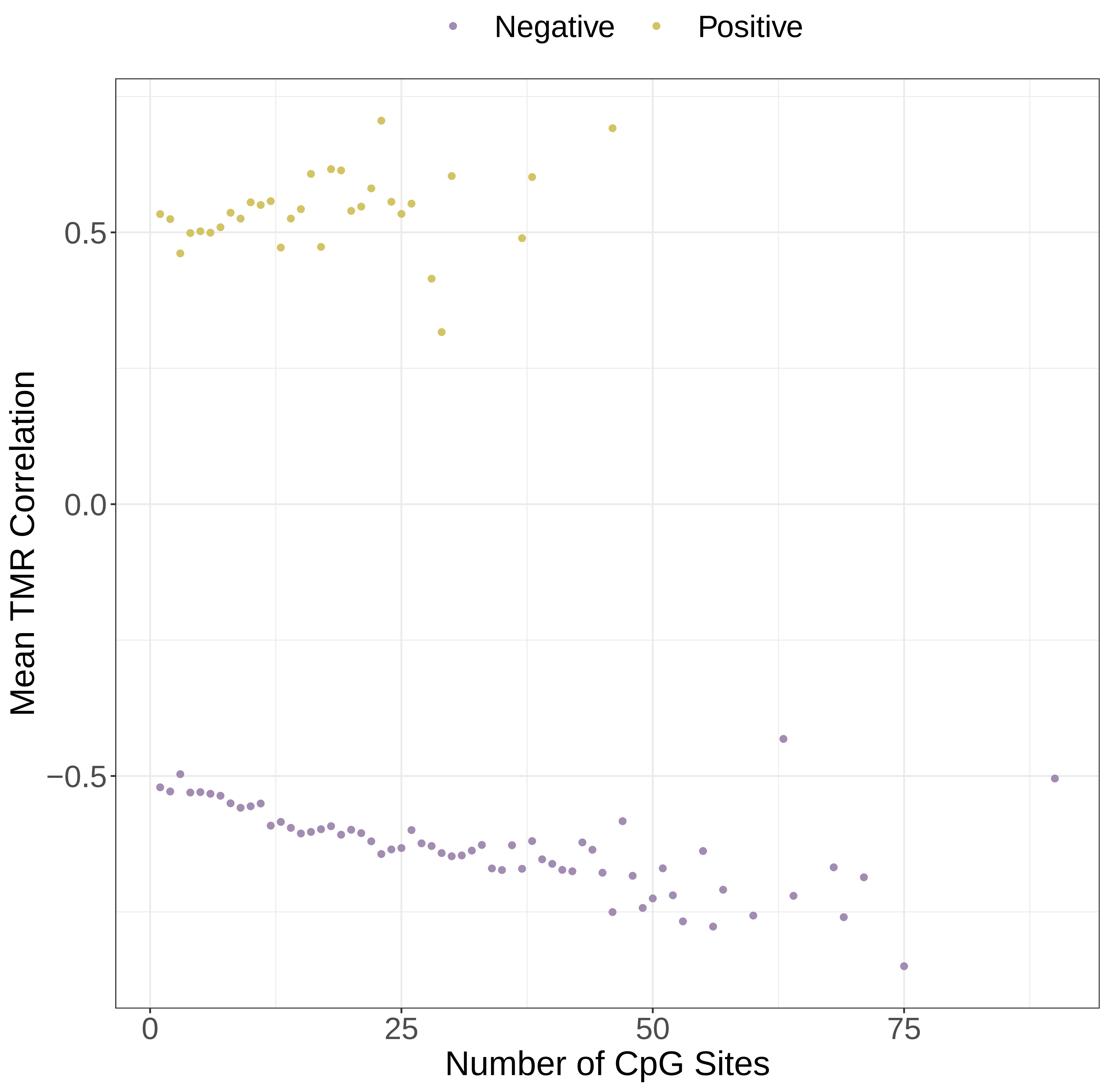
Negative
Positive
Mean TMR Correlation
0.5
0.0
−0.5
0
25
50
75
Number of CpG Sites

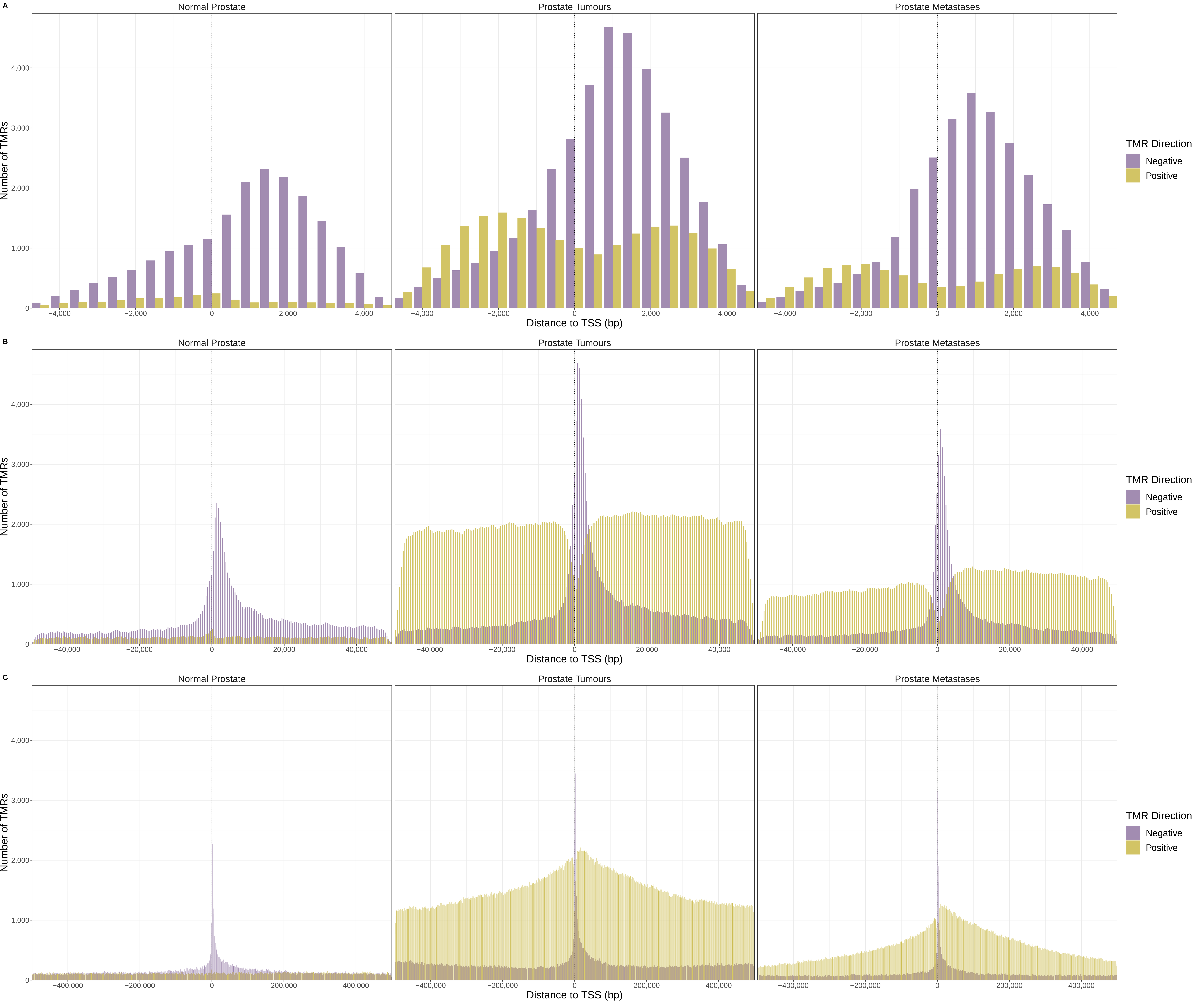

A
B
C
Normal Prostate
Prostate Tumours
Prostate Metastases
Number of TMRs
Distance to TSS (bp)
TMR Direction
Negative
Positive

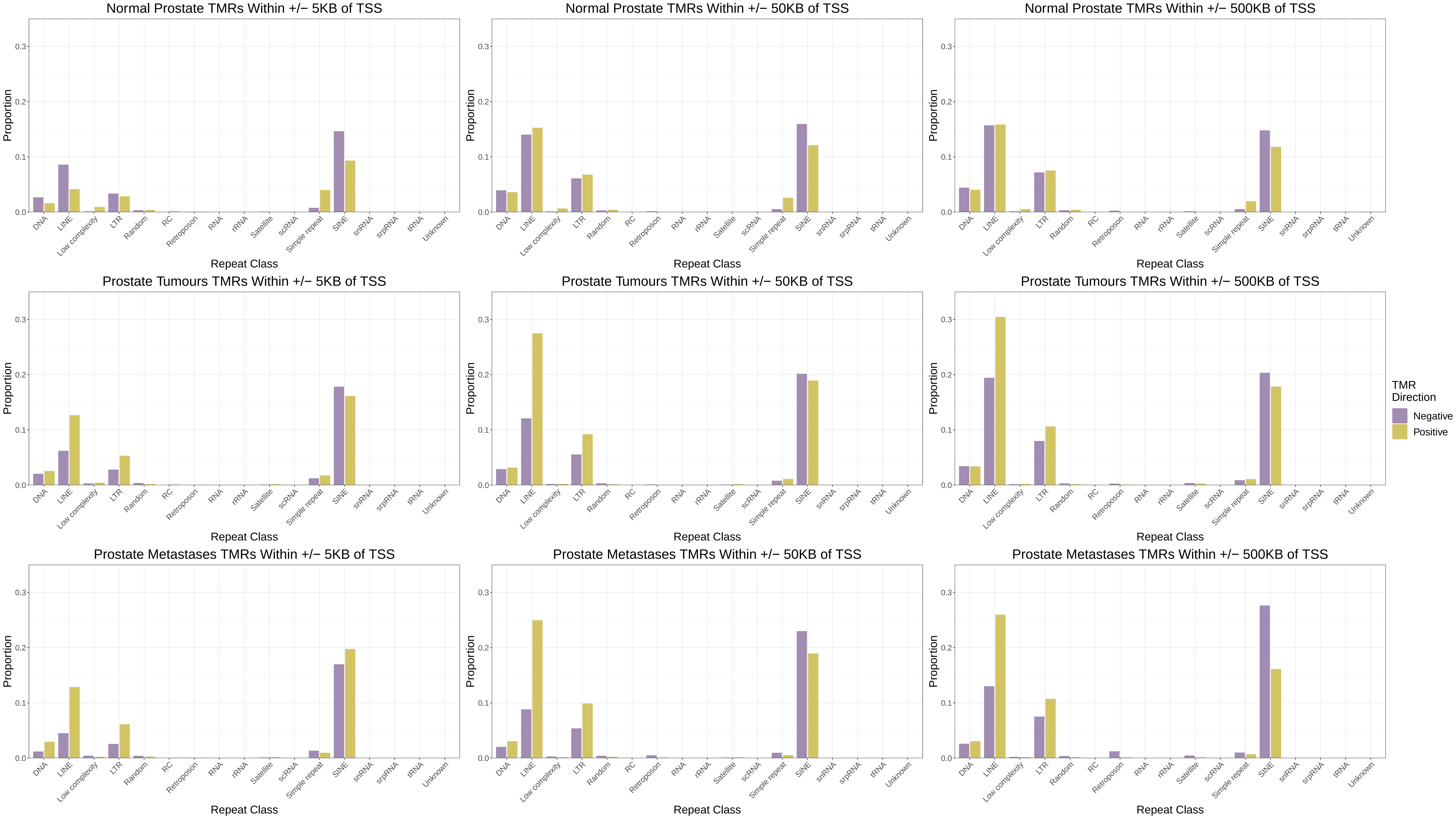

Normal Prostate TMRs Within +/− 5KB of TSS
Normal Prostate TMRs Within +/− 50KB of TSS
Normal Prostate TMRs Within +/− 500KB of TSS
Prostate Tumours TMRs Within +/− 5KB of TSS
Prostate Tumours TMRs Within +/− 50KB of TSS
Prostate Tumours TMRs Within +/− 500KB of TSS
Prostate Metastases TMRs Within +/− 5KB of TSS
Prostate Metastases TMRs Within +/− 50KB of TSS
Prostate Metastases TMRs Within +/− 500KB of TSS
Proportion
0.0
0.1
0.2
0.3
Repeat Class
DNA
LINE
Low complexity
LTR
Random
RC
Retroposon
RNA
rRNA
Satellite
scRNA
Simple repeat
SINE
snRNA
srpRNA
tRNA
Unknown
TMR Direction
Negative
Positive

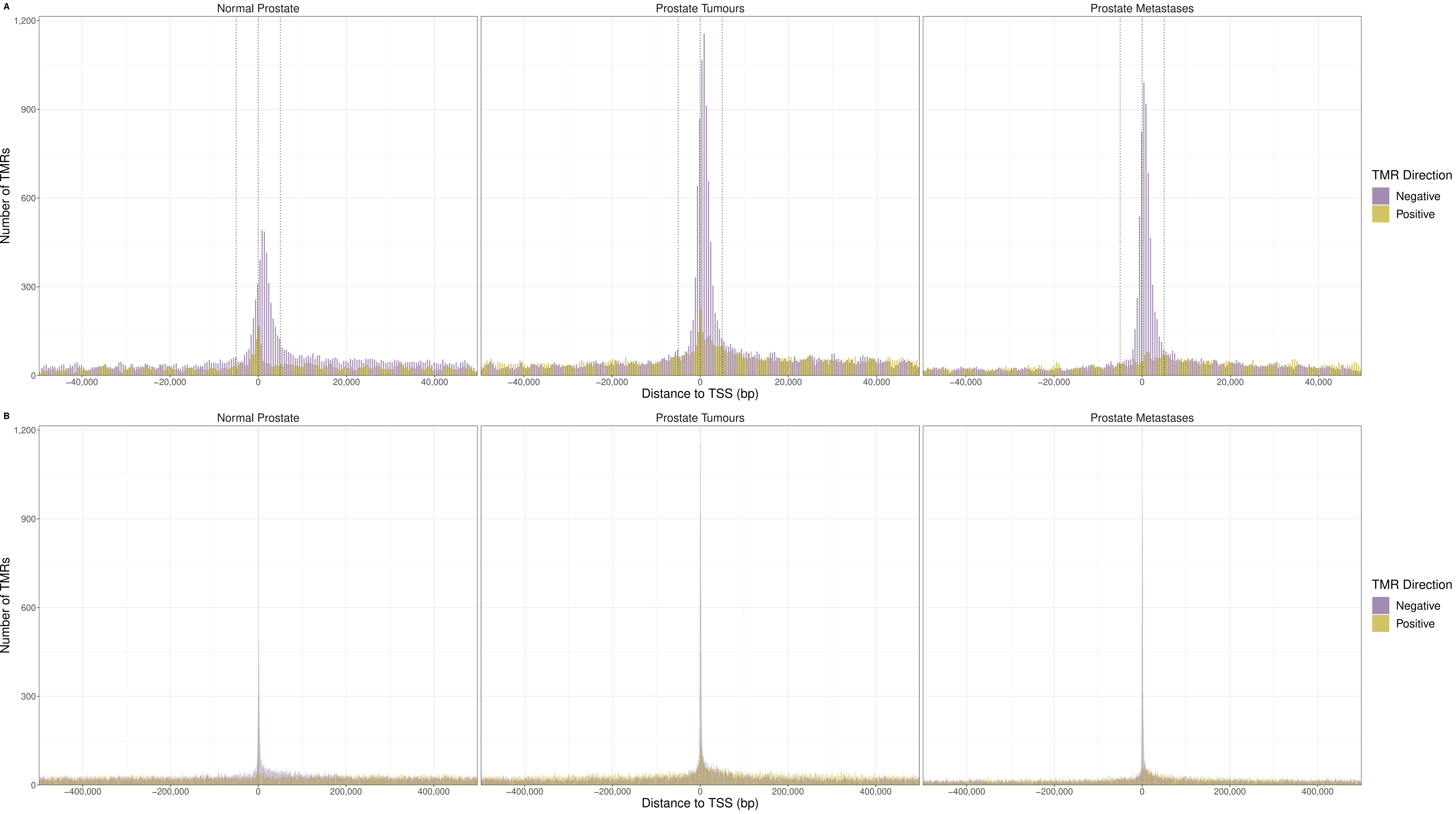
A
Normal Prostate
Prostate Tumours
Prostate Metastases
Number of TMRs
1,200
900
600
300
0
−40,000
−20,000
0
20,000
40,000
Distance to TSS (bp)
TMR Direction
Negative
Positive
B
Normal Prostate
Prostate Tumours
Prostate Metastases
Number of TMRs
−400,000
−200,000
0
200,000
400,000
Distance to TSS (bp)
TMR Direction
Negative
Positive

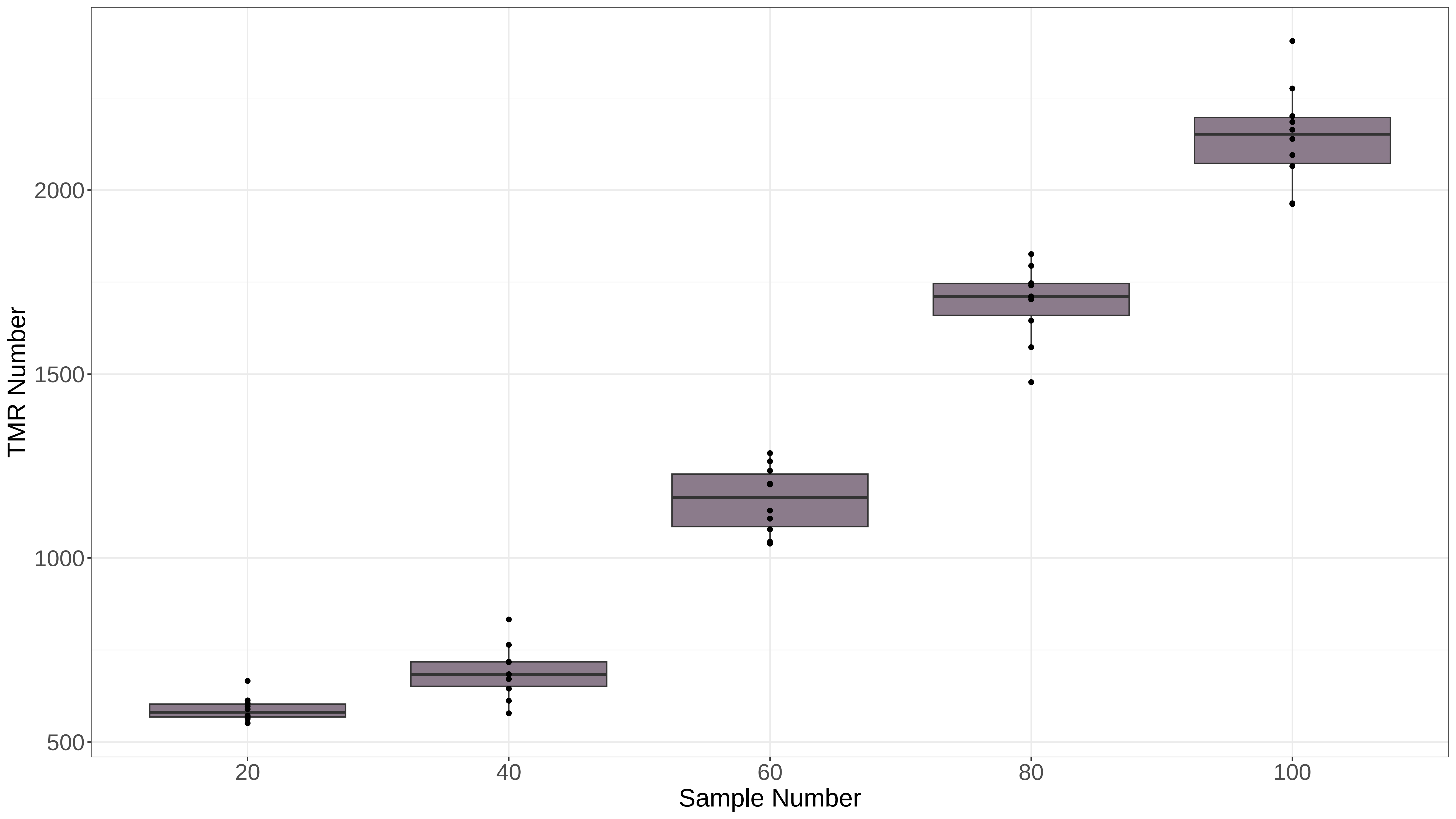

TMR Number
2000
1500
1000
500
20
40
60
80
100
Sample Number

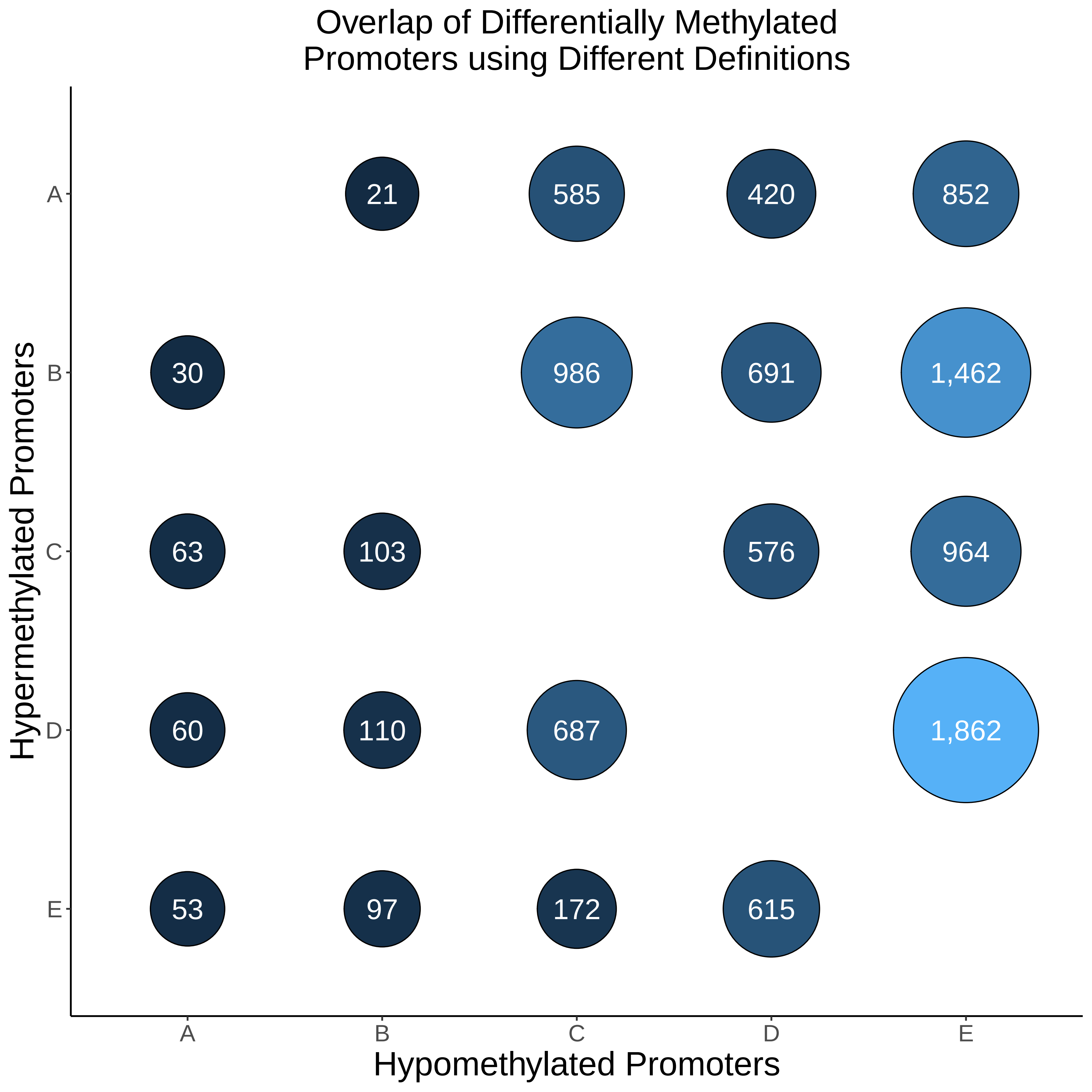

Overlap of Differentially Methylated
Promoters using Different Definitions
Hypermethylated Promoters
Hypomethylated Promoters
A
B
C
D
E
21
585
420
852
30
986
691
1,462
63
103
576
964
60
110
687
1,862
53
97
172
615

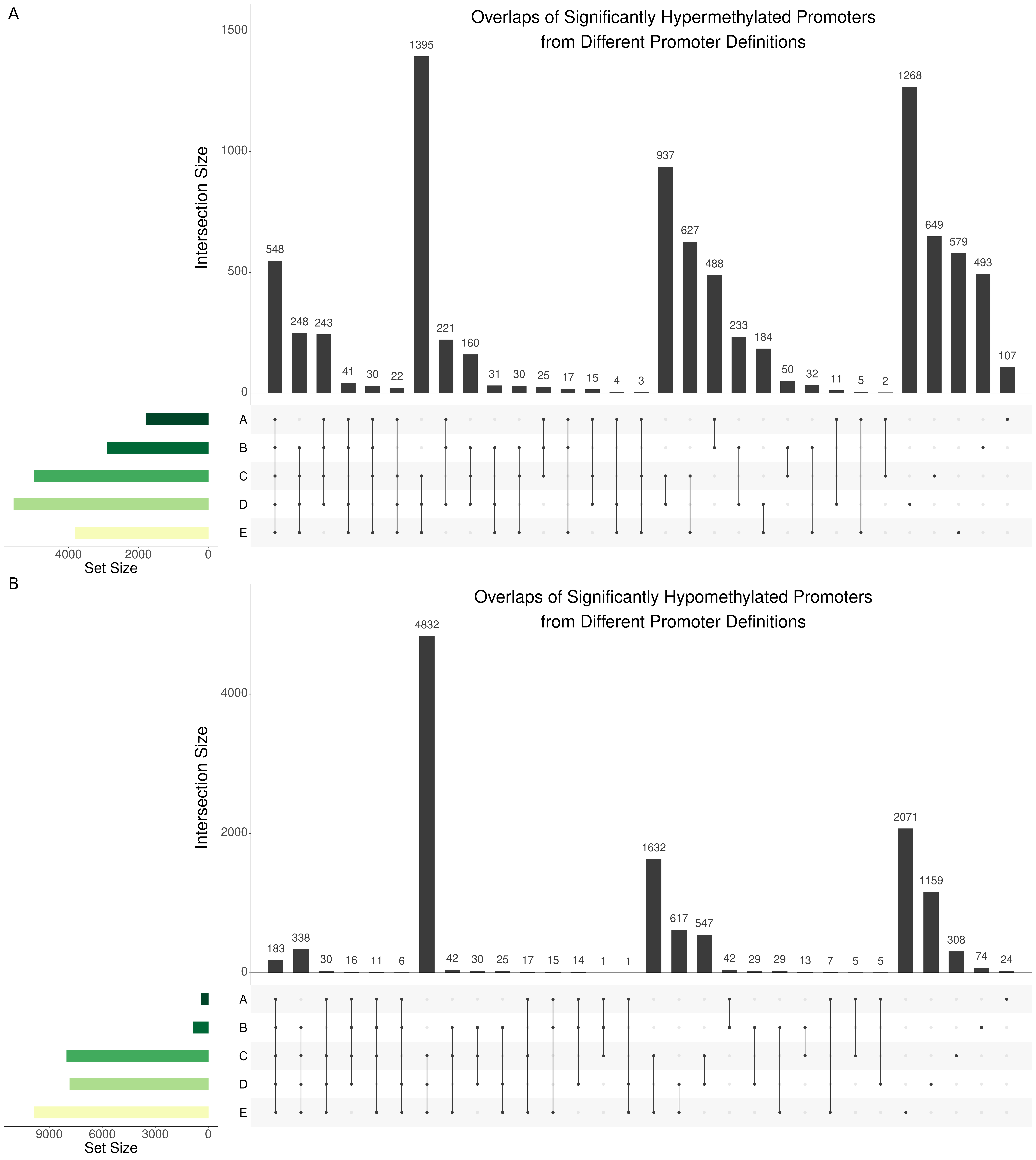
A
Overlaps of Significantly Hypermethylated Promoters
from Different Promoter Definitions
Intersection Size
1500
1000
500
0
548
248
243
41
30
22
1395
221
160
31
30
25
17
15
4
3
937
627
488
233
184
50
32
11
5
2
1268
649
579
493
107
A
B
C
D
E
4000
2000
0
Set Size
B
Overlaps of Significantly Hypomethylated Promoters
from Different Promoter Definitions
Intersection Size
4000
2000
0
183
338
30
16
11
6
4832
42
30
25
17
15
14
1
1
1632
617
547
42
29
29
13
7
5
5
2071
1159
308
74
24
A
B
C
D
E
9000
6000
3000
0
Set Size

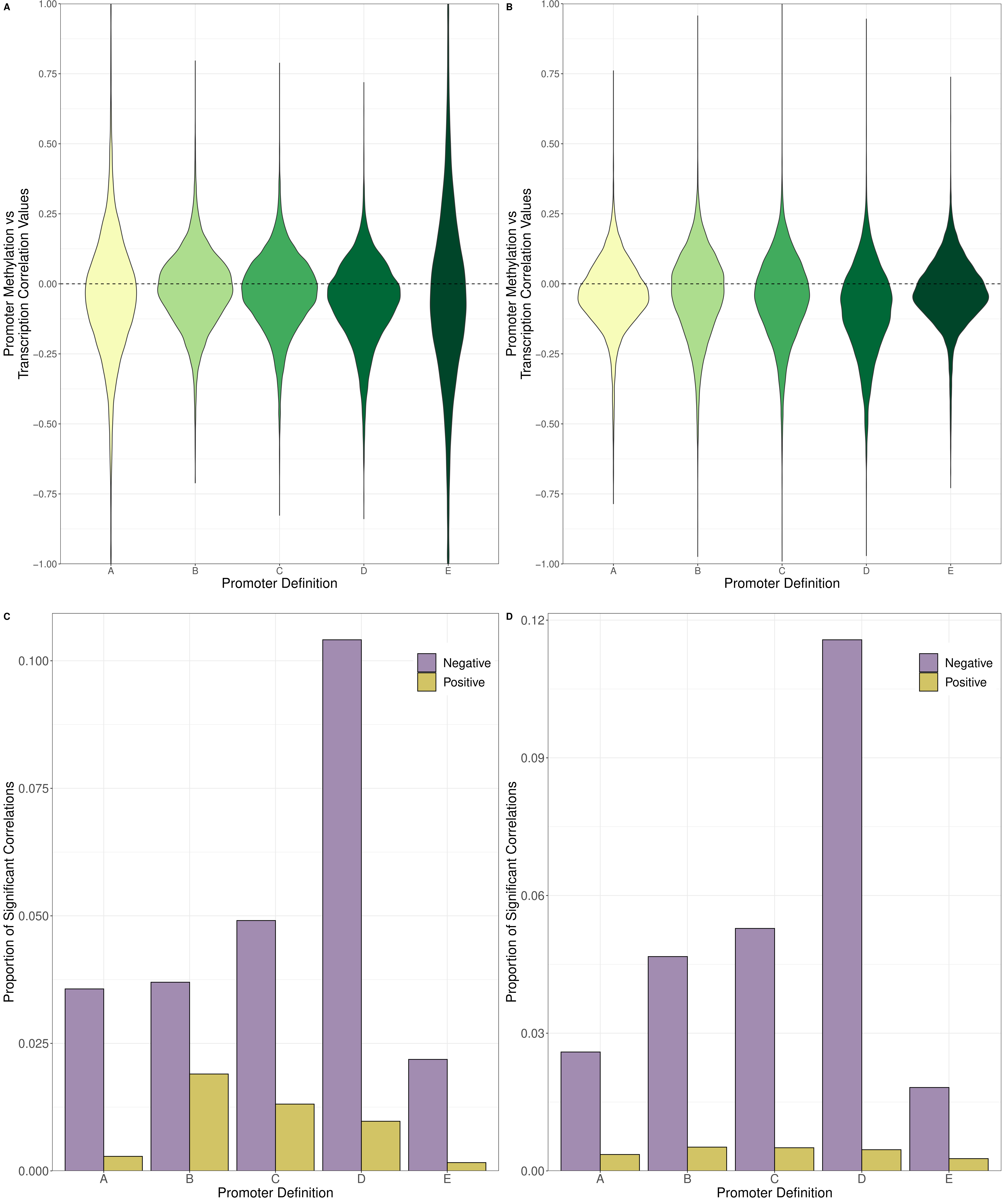

A
B
C
D
Promoter Methylation vs Transcription Correlation Values
Promoter Definition
Proportion of Significant Correlations
Negative
Positive
1.00
0.75
0.50
0.25
0.00
−0.25
−0.50
−0.75
−1.00
0.100
0.075
0.050
0.025
0.000
0.12
0.09
0.06
0.03
0.00
A
B
C
D
E

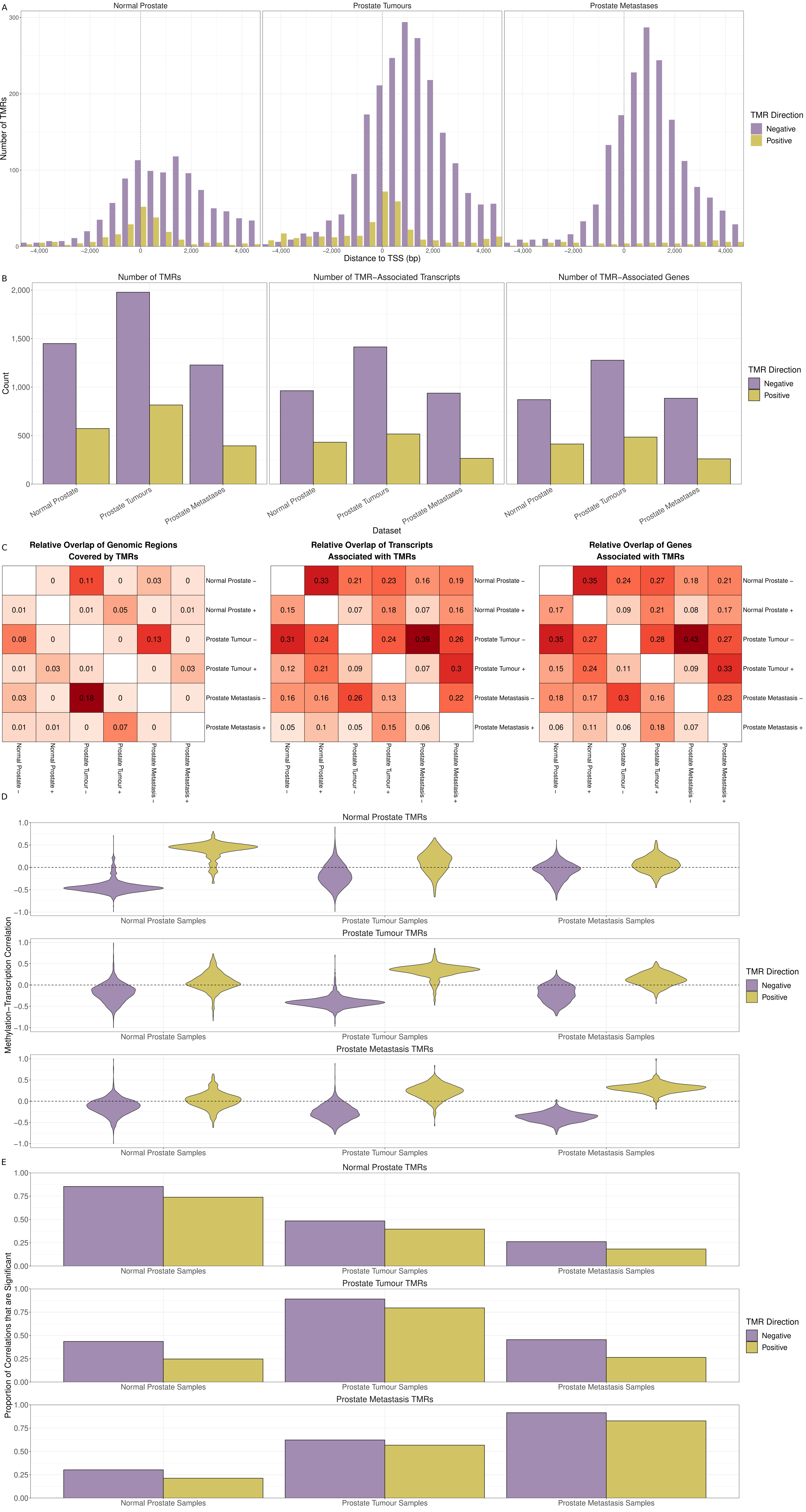
A
Normal Prostate
Prostate Tumours
Prostate Metastases
Number of TMRs
Distance to TSS (bp)
TMR Direction
Negative
Positive
B
Number of TMRs
Number of TMR−Associated Transcripts
Number of TMR−Associated Genes
Count
Dataset
Normal Prostate
Prostate Tumours
Prostate Metastases
C
Relative Overlap of Genomic Regions Covered by TMRs
Relative Overlap of Transcripts Associated with TMRs
Relative Overlap of Genes Associated with TMRs
Normal Prostate −
Normal Prostate +
Prostate Tumour −
Prostate Tumour +
Prostate Metastasis −
Prostate Metastasis +
0 0.11 0 0.03 0
0.01 0.01 0.05 0 0.01
0.08 0 0 0.13 0
0.01 0.03 0.01 0 0.03
0.03 0 0.18 0 0
0.01 0.01 0 0.07 0
0.33 0.21 0.23 0.16 0.19
0.15 0.07 0.18 0.07 0.16
0.31 0.24 0.24 0.39 0.26
0.12 0.21 0.09 0.07 0.3
0.16 0.16 0.26 0.13 0.22
0.05 0.1 0.05 0.15 0.06
0.35 0.24 0.27 0.18 0.21
0.17 0.09 0.21 0.08 0.17
0.35 0.27 0.28 0.43 0.27
0.15 0.24 0.11 0.09 0.33
0.18 0.17 0.3 0.16 0.23
0.06 0.11 0.06 0.18 0.07
D
Normal Prostate TMRs
Prostate Tumour TMRs
Prostate Metastasis TMRs
Methylation−Transcription Correlation
Normal Prostate Samples
Prostate Tumour Samples
Prostate Metastasis Samples
E
Proportion of Correlations that are Significant

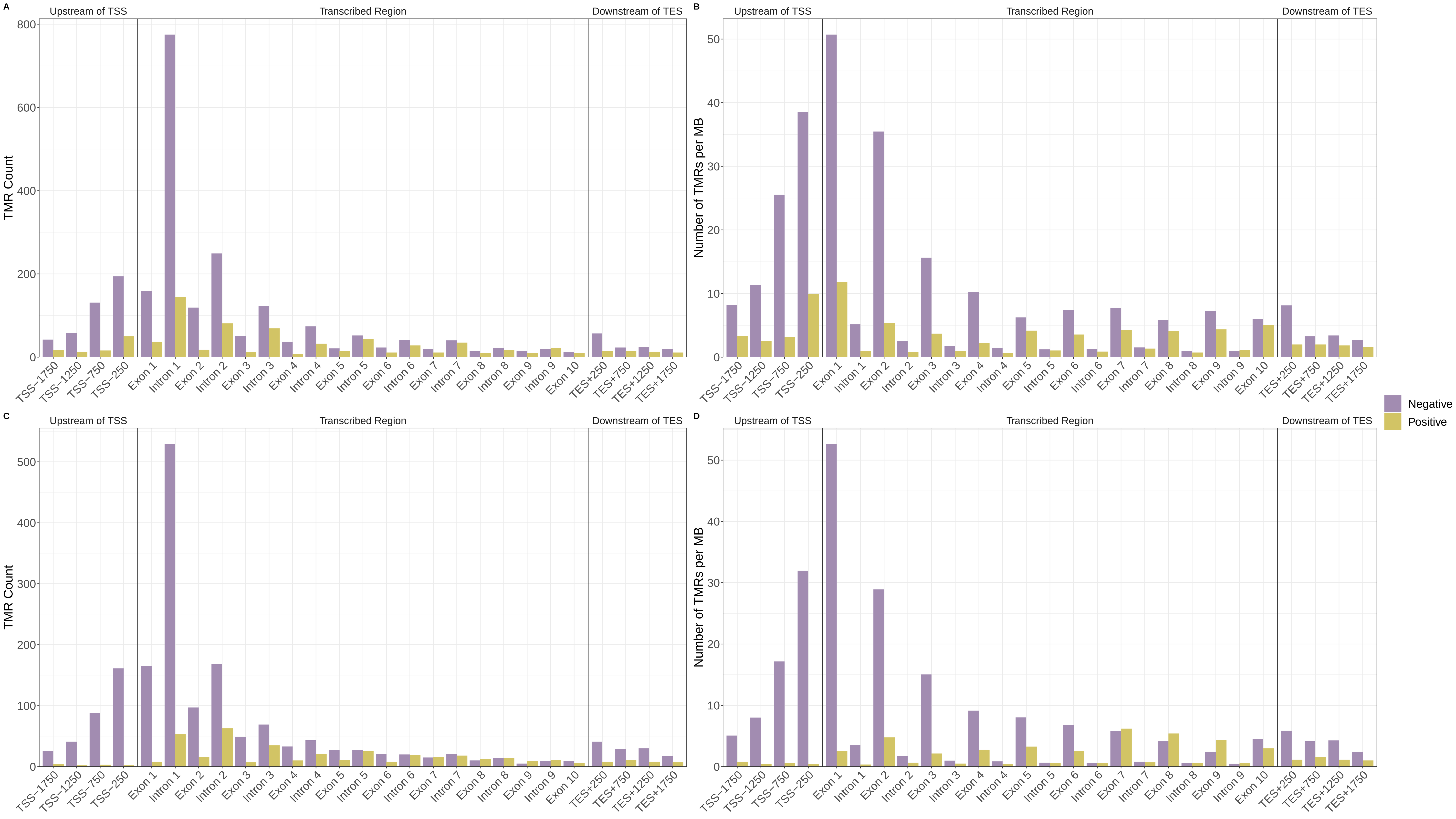
A
B
C
D
Upstream of TSS
Transcribed Region
Downstream of TES
TMR Count
Number of TMRs per MB
TSS−1750
TSS−1250
TSS−750
TSS−250
Exon 1
Intron 1
Exon 2
Intron 2
Exon 3
Intron 3
Exon 4
Intron 4
Exon 5
Intron 5
Exon 6
Intron 6
Exon 7
Intron 7
Exon 8
Intron 8
Exon 9
Intron 9
Exon 10
TES+250
TES+750
TES+1250
TES+1750
Negative
Positive

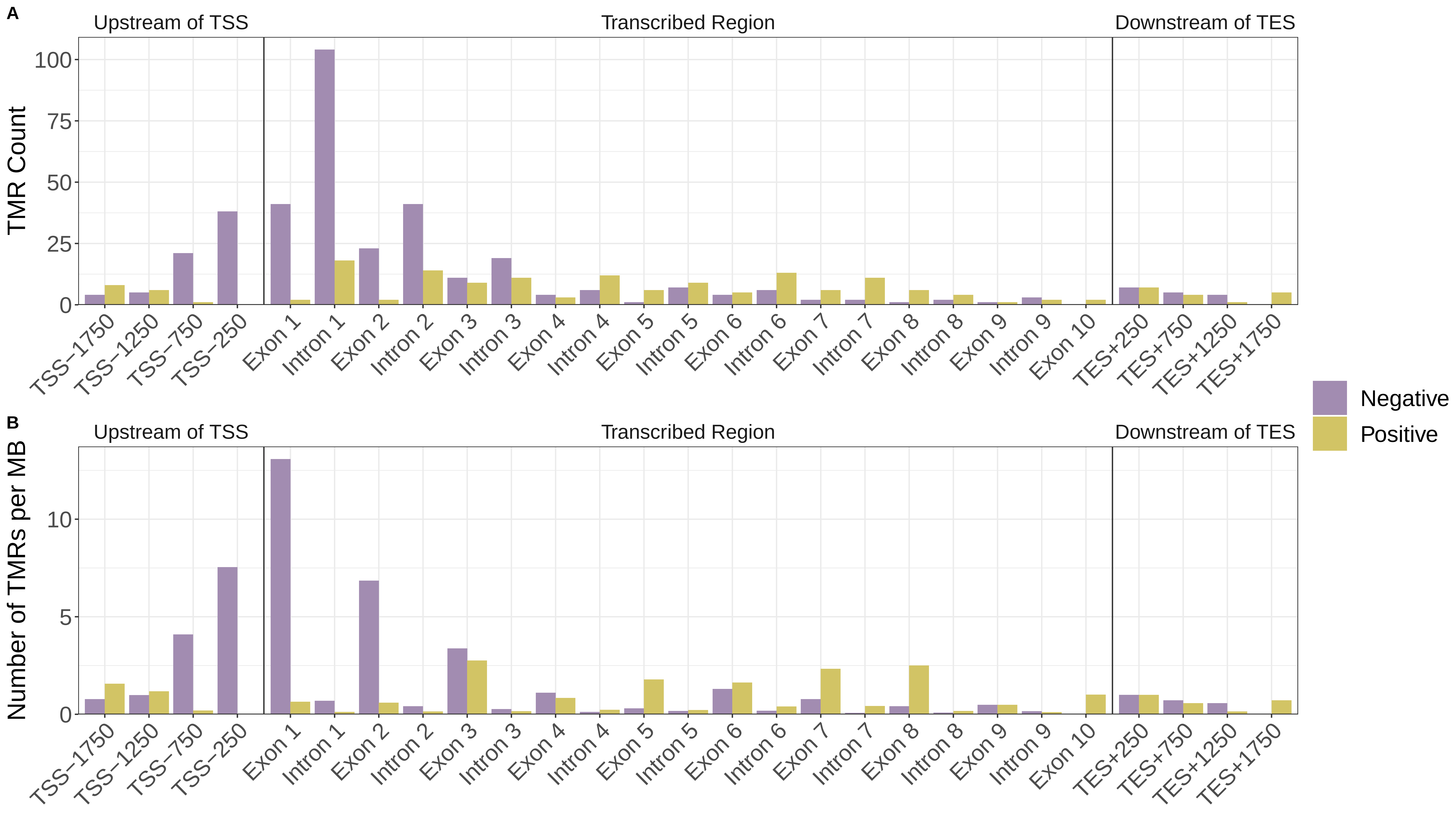
A
Upstream of TSS
Transcribed Region
Downstream of TES
TMR Count
100
75
50
25
0
TSS−1750
TSS−1250
TSS−750
TSS−250
Exon 1
Intron 1
Exon 2
Intron 2
Exon 3
Intron 3
Exon 4
Intron 4
Exon 5
Intron 5
Exon 6
Intron 6
Exon 7
Intron 7
Exon 8
Intron 8
Exon 9
Intron 9
Exon 10
TES+250
TES+750
TES+1250
TES+1750
Negative
Positive
B
Upstream of TSS
Transcribed Region
Downstream of TES
Number of TMRs per MB
10
5
0
TSS−1750
TSS−1250
TSS−750
TSS−250
Exon 1
Intron 1
Exon 2
Intron 2
Exon 3
Intron 3
Exon 4
Intron 4
Exon 5
Intron 5
Exon 6
Intron 6
Exon 7
Intron 7
Exon 8
Intron 8
Exon 9
Intron 9
Exon 10
TES+250
TES+750
TES+1250
TES+1750

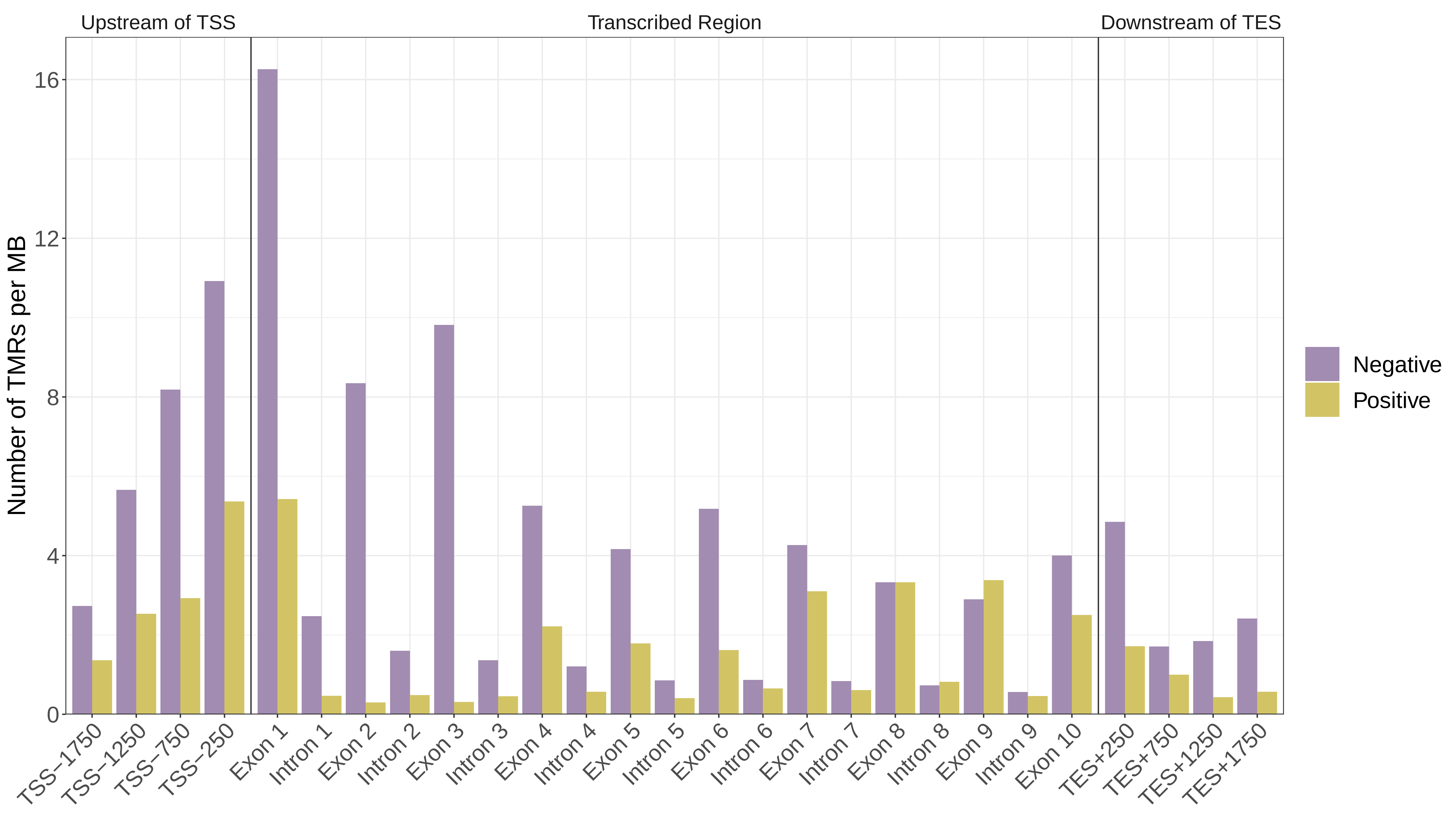

Upstream of TSS
Transcribed Region
Downstream of TES
Number of TMRs per MB
0
4
8
12
16
TSS−1750
TSS−1250
TSS−750
TSS−250
Exon 1
Intron 1
Exon 2
Intron 2
Exon 3
Intron 3
Exon 4
Intron 4
Exon 5
Intron 5
Exon 6
Intron 6
Exon 7
Intron 7
Exon 8
Intron 8
Exon 9
Intron 9
Exon 10
TES+250
TES+750
TES+1250
TES+1750
Negative
Positive

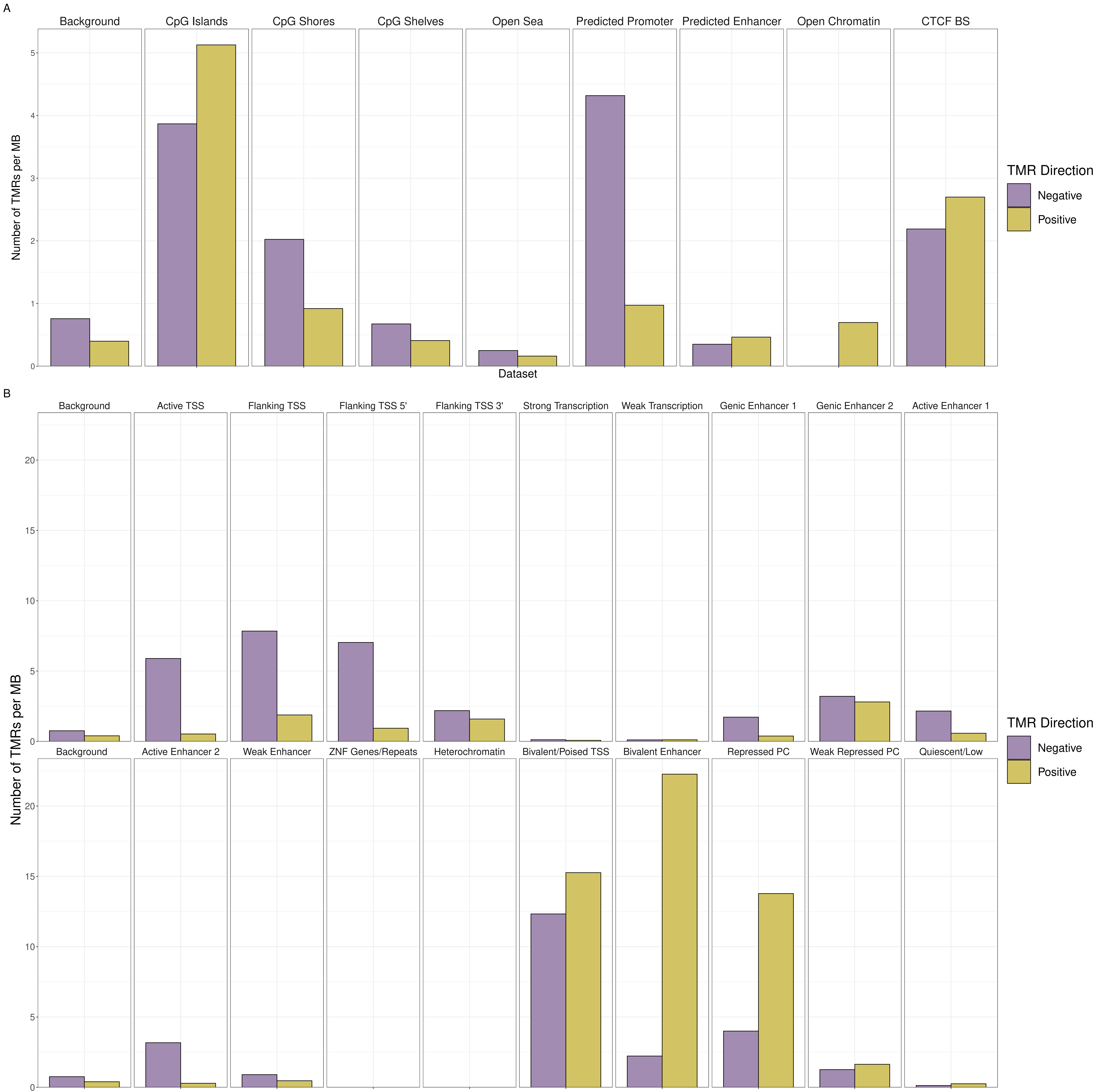
A
Background
CpG Islands
CpG Shores
CpG Shelves
Open Sea
Predicted Promoter
Predicted Enhancer
Open Chromatin
CTCF BS
Number of TMRs per MB
Dataset
TMR Direction
Negative
Positive
B
Background
Active TSS
Flanking TSS
Flanking TSS 5'
Flanking TSS 3'
Strong Transcription
Weak Transcription
Genic Enhancer 1
Genic Enhancer 2
Active Enhancer 1
Background
Active Enhancer 2
Weak Enhancer
ZNF Genes/Repeats
Heterochromatin
Bivalent/Poised TSS
Bivalent Enhancer
Repressed PC
Weak Repressed PC
Quiescent/Low
Number of TMRs per MB
TMR Direction
Negative
Positive

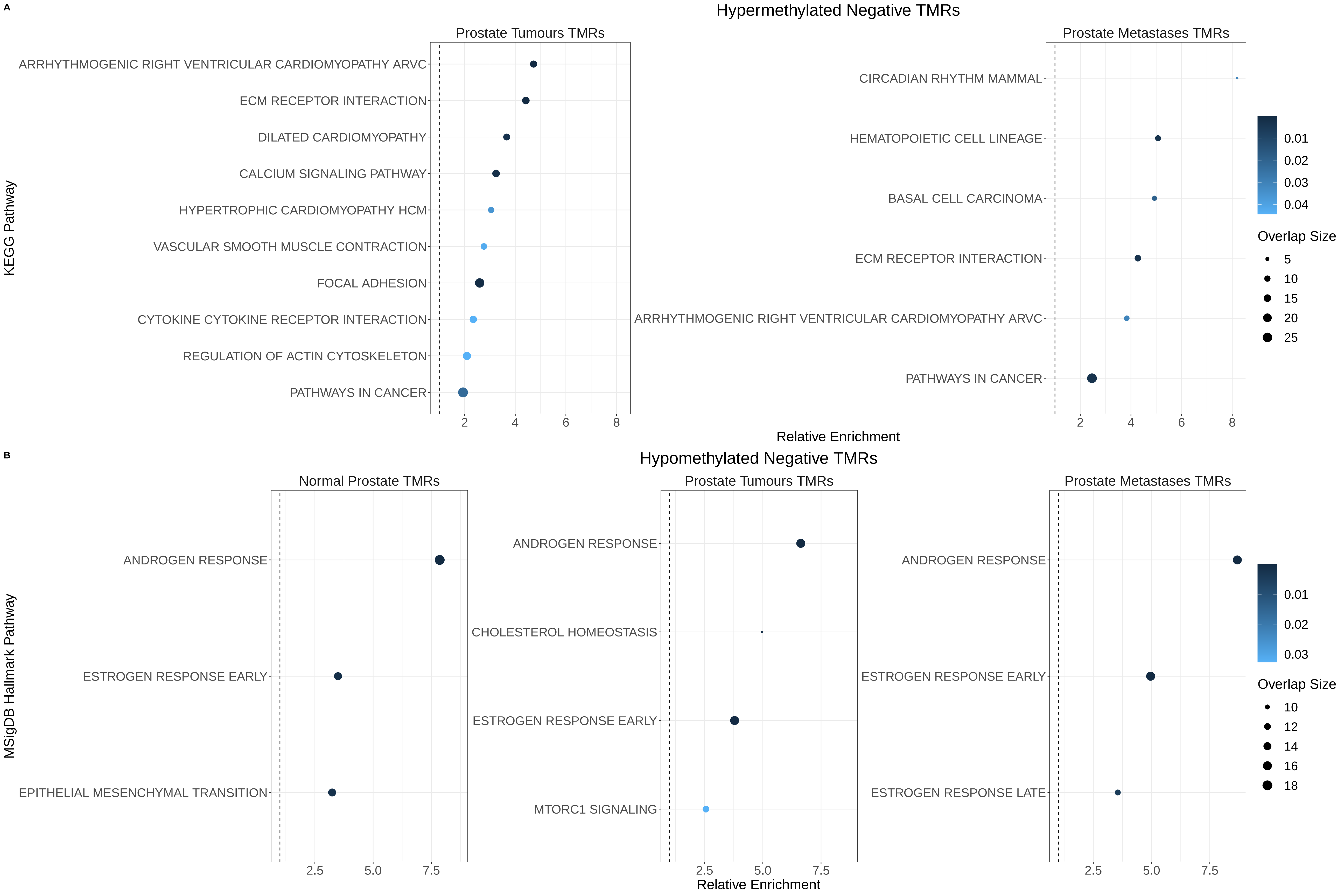
A
Hypermethylated Negative TMRs
Prostate Tumours TMRs
ARRHYTHMOGENIC RIGHT VENTRICULAR CARDIOMYOPATHY ARVC
ECM RECEPTOR INTERACTION
DILATED CARDIOMYOPATHY
CALCIUM SIGNALING PATHWAY
HYPERTROPHIC CARDIOMYOPATHY HCM
VASCULAR SMOOTH MUSCLE CONTRACTION
FOCAL ADHESION
CYTOKINE CYTOKINE RECEPTOR INTERACTION
REGULATION OF ACTIN CYTOSKELETON
PATHWAYS IN CANCER
Prostate Metastases TMRs
CIRCADIAN RHYTHM MAMMAL
HEMATOPOIETIC CELL LINEAGE
BASAL CELL CARCINOMA
ECM RECEPTOR INTERACTION
ARRHYTHMOGENIC RIGHT VENTRICULAR CARDIOMYOPATHY ARVC
PATHWAYS IN CANCER
KEGG Pathway
2 4 6 8
Relative Enrichment
0.01 0.02 0.03 0.04
Overlap Size
5 10 15 20 25
B
Hypomethylated Negative TMRs
Normal Prostate TMRs
ANDROGEN RESPONSE
ESTROGEN RESPONSE EARLY
EPITHELIAL MESENCHYMAL TRANSITION
Prostate Tumours TMRs
ANDROGEN RESPONSE
CHOLESTEROL HOMEOSTASIS
ESTROGEN RESPONSE EARLY
MTORC1 SIGNALING
Prostate Metastases TMRs
ANDROGEN RESPONSE
ESTROGEN RESPONSE EARLY
ESTROGEN RESPONSE LATE
MSigDB Hallmark Pathway
2.5 5.0 7.5
Relative Enrichment
0.01 0.02 0.03
Overlap Size
10 12 14 16 18

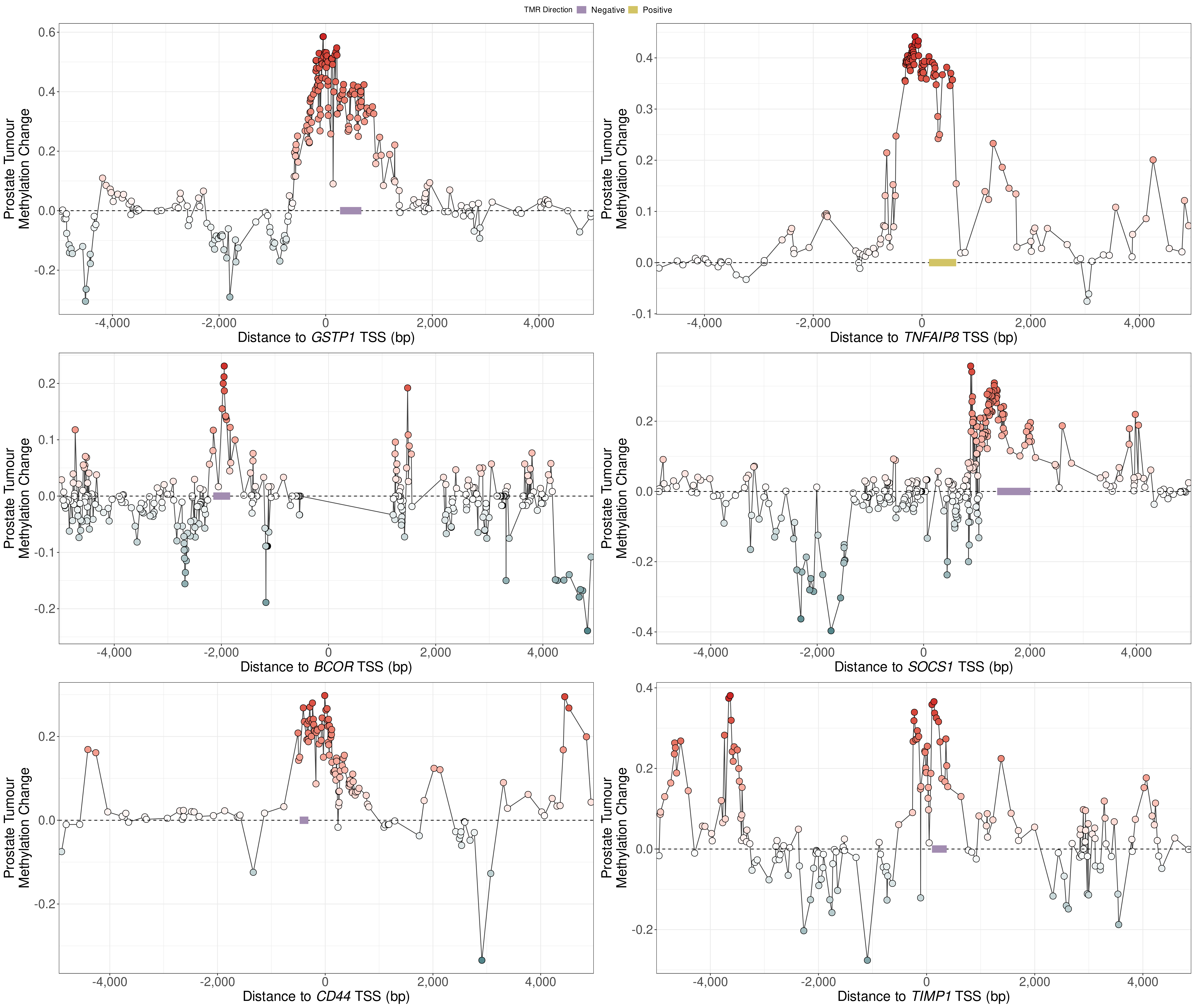

TMR Direction
Negative
Positive
Prostate Tumour Methylation Change
Distance to GSTP1 TSS (bp)
Distance to TNFAIP8 TSS (bp)
Distance to BCOR TSS (bp)
Distance to SOCS1 TSS (bp)
Distance to CD44 TSS (bp)
Distance to TIMP1 TSS (bp)

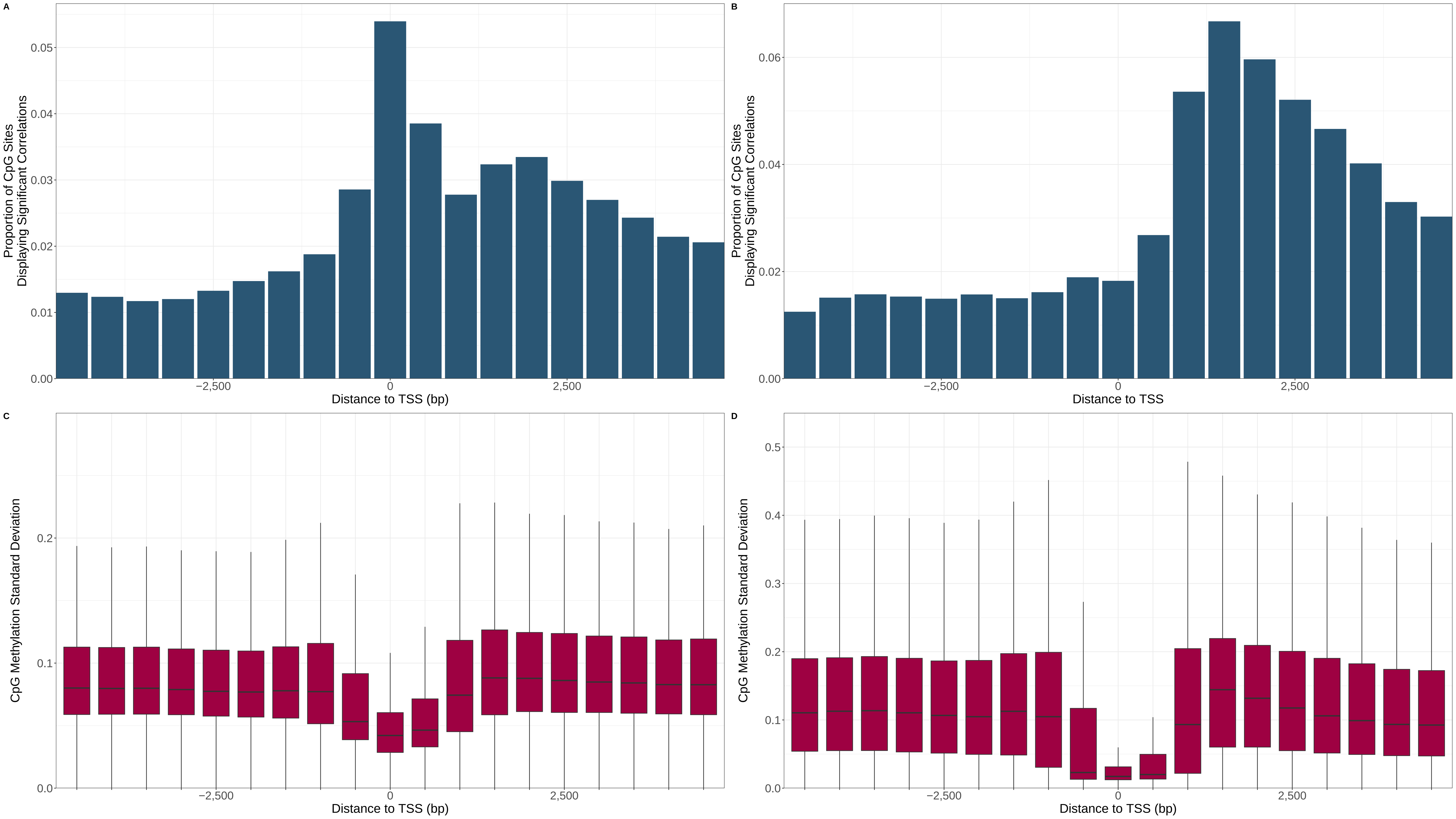

A
Proportion of CpG Sites Displaying Significant Correlations
0.00
0.01
0.02
0.03
0.04
0.05
−2,500
0
2,500
Distance to TSS (bp)
B
Proportion of CpG Sites Displaying Significant Correlations
0.00
0.02
0.04
0.06
−2,500
0
2,500
Distance to TSS
C
CpG Methylation Standard Deviation
0.0
0.1
0.2
−2,500
0
2,500
Distance to TSS (bp)
D
CpG Methylation Standard Deviation
0.0
0.1
0.2
0.3
0.4
0.5
−2,500
0
2,500
Distance to TSS (bp)

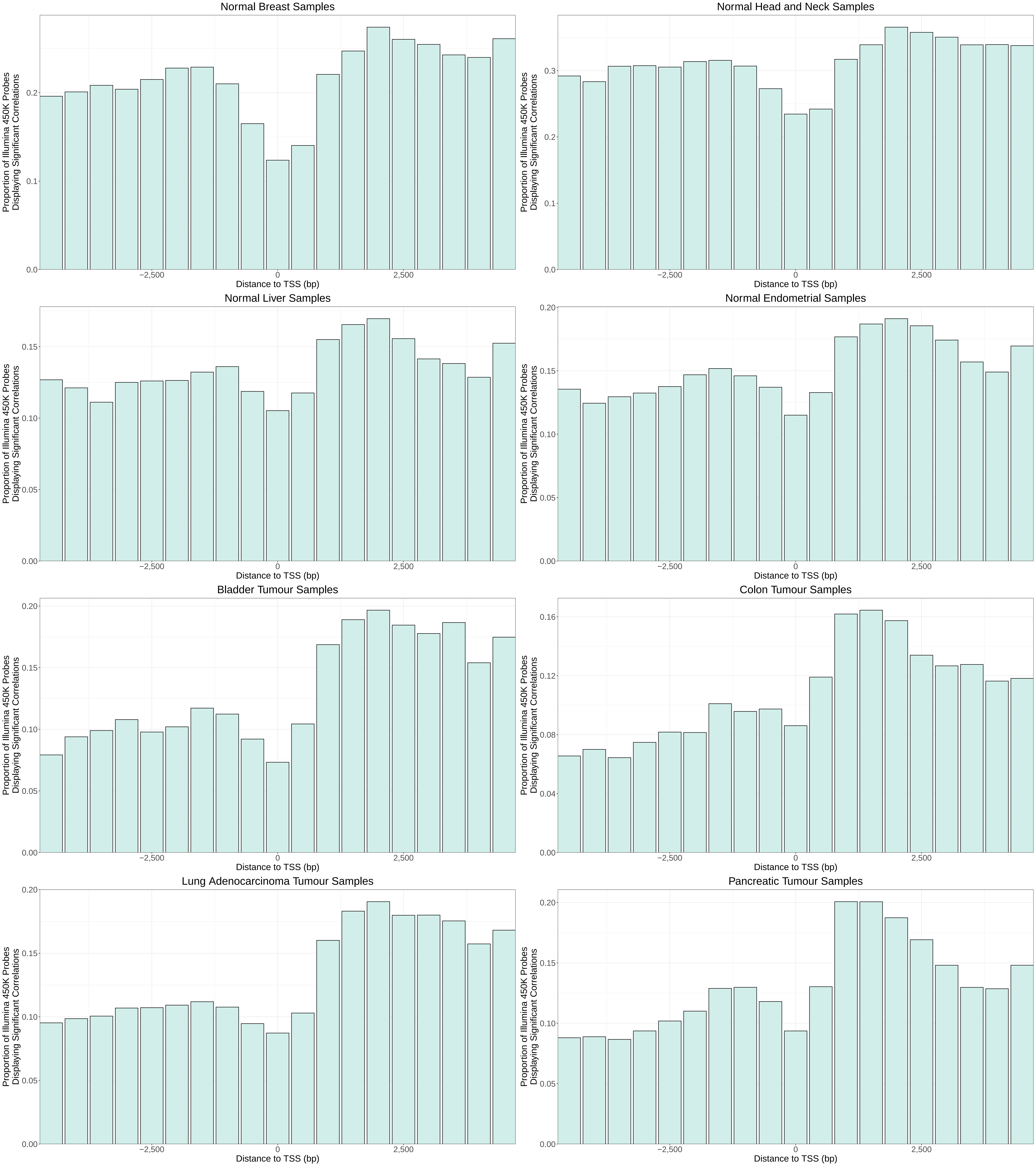
Normal Breast Samples
Normal Head and Neck Samples
Normal Liver Samples
Normal Endometrial Samples
Bladder Tumour Samples
Colon Tumour Samples
Lung Adenocarcinoma Tumour Samples
Pancreatic Tumour Samples
Proportion of Illumina 450K Probes Displaying Significant Correlations
Distance to TSS (bp)
−2,500
0
2,500

Supplementary Figure Legends

**Supplementary Figure 1: (A)** The number of TMRs found in prostate metastases using different combinations of the number of flanking CpGs used to construct windows and the smoothing factor used for the exponential moving average. **(B)** The proportion of TMRs identified in prostate metastases using different parameter combinations resulting in significant correlations when evaluating their correlations in prostate tumour samples. 10 flanking CpGs and a smoothing factor of 0.75 seemed to give a good trade-off between the number of TMRs discovered and the proportion of these that gave significant correlations.

**Supplementary Figure 2:** The mean methylation-transcription correlations for TMRs identified in normal prostate samples grouped by the number of CpG sites that they overlap. There was a Spearman correlation value of 0.3 (p-value < 2.2e-16) between the number of CpG sites and the absolute methylation-transcription correlations of TMRs.

**Supplementary Figure 3:** The relative location of negative and positive TMRs in normal prostate, prostate tumours and prostate metastases within +/- 5 KB (A), +/- 50 KB (B) and +/- 500 KB (C) of the TSS without removal of TMRs overlapping repeat elements. The regions around TSS were divided into 500 bp bins and the number of times TMRs overlapped these bins were counted. The x-axis shows the distance from the center of bins to the TSS. The dotted line indicates the location of the TSS. There are a huge number of positive TMRs found surrounding the TSS in prostate tumour and metastasis samples.

**Supplementary Figure 4:** Overlap of TMRs with repeat elements. X-axis shows the class of repetitive element and Y-axis shows the proportion of TMRs overlapping elements of the indicated class. TMRs which had at least 25% of their sequence overlapping repeats from the indicated class were defined as overlapping that class. Top row shows overlaps for TMRs identified in normal prostate samples, the middle row shows overlaps for TMRs identified in prostate tumour samples and bottom row shows overlaps for TMRs identified in prostate metastasis samples, each within 5 kb, 50 kb and 500 kb of the TSS. The proportion of TMRs from prostate tumours and prostate metastases overlapping certain classes of repeats, particularly that of positive TMRs with LINE repeats, generally increases as the search region is expanded.

**Supplementary Figure 5:** Distribution of TMRs (A) 50 KB and (B) 500 KB upstream and downstream of TSS after removal of TMRs overlapping repeats. Axes and colours are as in Supplementary Figure 3. Dotted lines in top panels show the location of -5 kb and + 5 kb. Compared to Supplementary Figure 3 which showed the distribution of TMRs before removal of repeats, there are relatively few TMRs located beyond 5 kb from the TSS. The region just downstream of the TSS remains the most common location for negative TMRs.

**Supplementary Figure 6:** The number of TMRs identified in prostate tumour samples with different sized subsets of samples. 10 random samples were used for each sample size. Below about 60 samples, the number of TMRs identified is relatively small.

**Supplementary Figure 7:** The intersection sizes of hypermethylated promoters identified using one of five different promoter definitions with hypomethylated promoters identified using another one of the definitions.

**Supplementary Figure 8:** UpSet plots showing the intersection of hypermethylated (A) and hypomethylated (B) promoters resulting from the use of different promoter definitions.

**Supplementary Figure 9:** The distribution of promoter methylation-transcription Spearman correlation values for all protein-coding transcripts using each of the 5 different promoter definitions in prostate tumour samples **(A)** and prostate metastasis samples **(B)**. The proportion of statistically significant correlations for each promoter definition divided into negative and positive correlations in prostate tumour samples **(C)** and in prostate metastasis samples **(D)**.

**Supplementary Figure 10: (A)** Location of TMRs within +/- 5 kb of the TSS. Regions were divided into 500 bp bins and the number of times TMRs overlapped these bins is displayed in normal prostate samples, prostate tumour samples and prostate metastasis samples. The x-axis shows the distance from the center of bins to the TSS. The dotted lines indicate 5 kb upstream, the TSS (0) and 5 kb downstream. **(B)** The number of negative and positive TMRs discovered within +/- 5KB of TSS and numbers of different transcripts and genes associated with TMRs in normal prostate, prostate tumour samples and prostate metastasis samples. **(C)** Heatmaps of the relative overlaps of TMRs and TMR-associated genes and transcripts from normal prostate, prostate tumour samples and prostate metastasis samples. For TMRs, a TMR from one group was defined as overlapping TMRs from another group if at least 25% of its sequence was located in TMRs from the other group. The relative overlap of one group with another was then defined as the proportion of TMRs from the first group (corresponding to rows) which overlapped the second (corresponding to columns). For example, 33% of the negative TMRs identified in the prostate metastasis. **(D)** Distributions of correlation values of TMRs with their associated transcript. Each panel shows the correlation values for TMRs identified in one of the datasets in each of the three datasets: the dataset in which it was identified as well as the other two datasets. For example, the central plot in the bottom panel shows the correlation values between TMR methylation and expression of the associated transcript in prostate tumour samples for negative and positive TMRs identified in prostate metastasis samples. **(E)** Proportion of statistically significant correlations between methylation of TMRs and their associated transcript. Like in panel D, each panel represents the correlation values for TMRs identified in one of the datasets in each of the three datasets.

**Supplementary Figure 11: (A and B)** The number of TMRs identified in prostate tumour samples (A) and prostate metastasis samples (B) within transcribed regions or regions within 5 kb upstream of the TSS or 5 kb downstream of the TES. Transcribed regions were separated into individual introns and exons while upstream and downstream regions were divided into 500 bp bins with the x-axis label indicating the centre of the bins. **(C and D)** The number of TMRs per MB of the indicated class of region in prostate tumour samples (C) prostate metastasis samples (D). The first exon has the highest density of TMRs.

**Supplementary Figure 12: (A)** The number of TMRs identified in tissue samples from the Roadmap Epigenomics project within transcribed regions or regions within 5 kb upstream of the TSS or 5 kb downstream of the TES. Transcribed regions were separated into individual introns and exons while upstream and downstream regions were divided into 500 bp bins with the x-axis label indicating the centre of the bins. The greatest number of TMRs are found in the first intron. **(B)** The number of TMRs per MB of the indicated class of region. The first exon has the highest density of TMRs.

**Supplementary Figure 13:** The number of TMRs associated with the TSS for MANE transcripts identified per MB within transcribed regions or regions within 5 kb upstream of the TSS or 5 kb downstream of the TES in normal prostate samples. Transcribed regions were separated into individual introns and exons while upstream and downstream regions were divided into 500 bp bins with the x-axis label indicating the centre of the bins. The first exon has the greatest density of TMRs.

**Supplementary Figure 14: (A)** The normalized number of TMRs identified in samples from the Roadmap Epigenomics project overlapping different classes of genomic regulatory elements. Normalization was done to account for the fact that different classes of regulatory elements occupy varying proportions of the genome and was performed by dividing the number of TMRs overlapping each regulatory class by the number of MB of DNA in TMR search regions (the transcription units with 5 kb added upstream and downstream) overlapping the respective regulatory class. CTCF BS stands for CTCF binding site. **(B)** The normalized number of Roadmap TMRs overlapping different chromatin states, calculated similarly as with the genomic regulatory elements above.

**Supplementary Figure 15: (A)** Overrepresentation of KEGG pathways in genes associated with hypermethylated TMRs identified in prostate tumours and prostate metastasis samples. **(B)** Overrepresentation of MSigDB Hallmark pathways in genes associated with hypomethylated TMRs identified in normal prostate, prostate tumours and prostate metastasis samples. The androgen response is the top pathway for all three datasets.

**Supplementary Figure 16:** Methylation change at TMRs associated with *GSTP1*, *TNFAIP8*, *BCOR*, *SOCS1*, *CD44* or *TIMP1* (using the TSS associated with the transcripts ENST00000398606, ENST00000504771, ENST00000378444, ENST00000332029, ENST00000263398 and ENST00000456754) in prostate tumours. The x-axes show distance of CpG sites upstream and downstream of the TSS in base pairs and the y-axes show mean methylation change relative to normal prostate samples.

**Supplementary Figure 17: (A and B)** The proportion of CpG sites where DNA methylation was significantly correlated with transcription in 500 bp bins around the TSS in normal prostate samples (A) and prostate metastases (B). Significant was defined as a corrected p-value under 0.05 for the Spearman correlation values. **(C and D)** Standard deviation values for the methylation of CpG sites in 500 bp bins around TSS in normal prostate samples (C) and prostate metastases (D).

**Supplementary Figure 18:** The proportion of Illumina 450K probes where DNA methylation was significantly correlated with gene expression in 500 bp bins around TSS in normal and tumour samples from different cancer types from TCGA. The TSS associated with the MANE transcript was used as the TSS for each gene. Significant was defined as an FDR-corrected p-value $< 0.05$ for the Spearman correlation values. The greatest proportion of significant correlation is generally found downstream of the TSS, with the region at the TSS displaying the lowest proportion.

Supplementary Table legends

Supplementary Table 1: All significantly enriched transcriptional regulators for each TMRs group. The enrichment of binding sites among TMR groups was tested by comparing the proportion of CpG sites within TMRs which overlap binding sites with that of all CpGs within the regions +/- 5 kb around the associated TSS using a two-sided chi-squared test. p-values were adjusted using the Benjamini-Hochberg procedure.

Supplementary Table 2: File accession IDs and tissue types for files downloaded from the Roadmap Epigenomics Project.
